# Mental image reconstruction from human brain activity

**DOI:** 10.1101/2023.01.22.525062

**Authors:** Naoko Koide-Majima, Shinji Nishimoto, Kei Majima

**Author notes:** **Corresponding author**: Kei Majima, Ph.D. Institute for Quantum Life Science, National Institutes for Quantum Science and Technology, Chiba, 263-8555, Japan. These authors contributed equally to this work (Naoko Koide-Majima, Kei Majima). Declaration of interest: None.

## Abstract

Visual images perceived by humans can be reconstructed from their brain activity. However, the visualization (externalization) of mental imagery remains a challenge. In this study, we demonstrated that the visual image reconstruction method proposed in the seminal study by Shen et al. (2019) heavily relied on low-level visual information decoded from the brain and could not efficiently utilize semantic information that would be recruited during mental imagery. To address this limitation, we extended the previous method to a Bayesian estimation framework and introduced the assistance of semantic information into it. Our proposed framework successfully reconstructed both seen (i.e., directly captured by the human eye) and imagined images from the brain activity. These results suggest that our framework would provide a technology for directly investigating the subjective contents of the brain.

## Introduction

Neural decoding technologies enable the visualization of perceptual contents based on brain activity^1, 2^. Previous studies have demonstrated that images seen by human participants can be reconstructed from the brain activity measured using functional magnetic resonance imaging (fMRI). Several studies have reconstructed visual perception for specific domains such as faces^3, 4^, hand-written letters^5^, and binary images^6–8^. Other studies have decoded seen natural images^9, 10^ or videos^11^ using visual features inspired by neurophysiological discoveries. Recently, by incorporating the assistance of deep neural networks (DNNs)^12, 13^ and generative models^14–21^, several studies have achieved higher-fidelity natural image reconstruction^22–26^, which has become a tool for investigating the visual processing in the brain (e.g., visual representation, attention^27^, and illusion^28^).

Previous studies have succeeded in reconstructing images seen by humans from their brain activity; however, externalizing mental imagery remains a challenge. For example, in one of the previous studies, Shen et al. (2019)^23^ attempted to reconstruct both seen and imagined images, but the reconstruction of imagined images has remained rudimentary. As we will demonstrate, one possible reason is that this previous method heavily relied on low-level visual information decoded from the brain. According to other neuroimaging studies, high-level or semantic information (representation) is thought to be recruited more strongly in the brain during mental imagery than low-level visual information. Although low-level visual features of imagined images (e.g., Gabor-wavelet features) can be decoded to a certain extent^29–32^, high-level visual features are more helpful in identifying imagined objects from brain activity^33^. Furthermore, categories of imagined objects can be better predicted from the brain activity in high-level visual areas than in low-level visual areas^34–36^. Thus, high-level and semantic information should be efficiently incorporated into the image reconstruction method to successfully externalize mental imagery.

To overcome the limitations of Shen et al.’s (2019)^23^ method, we first extended this previous method to a Bayesian estimation framework and then introduced the assistance of semantic information. In the previous method^23^, brain activity measured by fMRI was first translated (decoded) into VGG19’s hierarchical representations^13^ (i.e., unit activations of individual layers in VGG19) using a variant of linear regression (Fig. 1a). Subsequently, an image was generated using an iterative process, such that the generated image would lead to unit activations similar to those decoded from the brain. The resulting image was considered a reconstruction. Whereas all convolutional and fully-connected layers of VGG19 were combined in the previous study, as we will demonstrate, this previous method failed to produce meaningful images using only high VGG layers. Accordingly, this method relied heavily on low-level visual information. In our current study, by viewing the image generation process in this method as maximum likelihood estimation, we extended it to Bayesian estimation (Fig. 1b). This framework enables us to use a sophisticated prior of natural images developed in recent computer vision studies, which is expected to help produce meaningful images even from abstract or partial information.

**Fig. 1.**
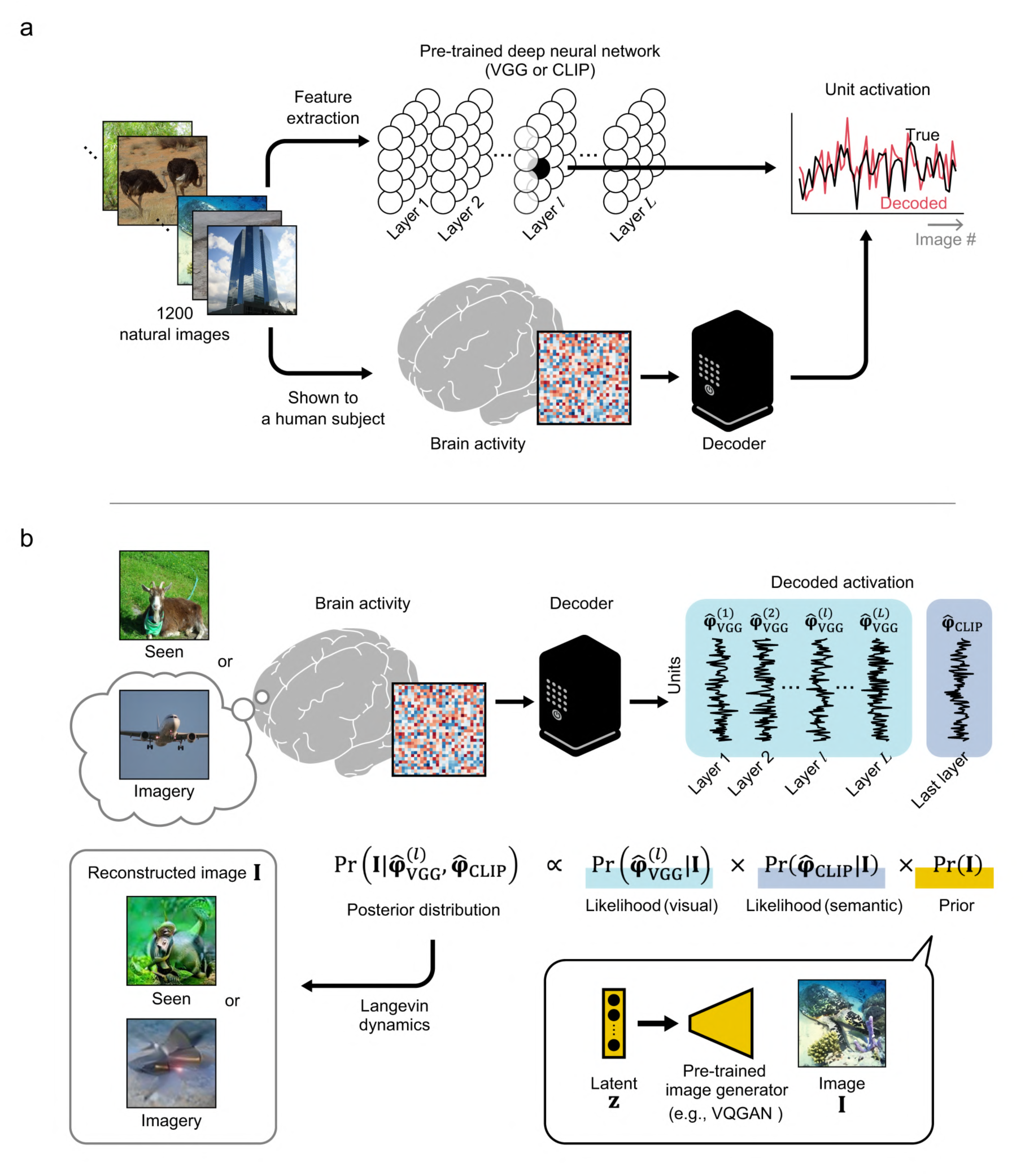
Proposed reconstruction framework. **(a)** Decoder training. In our framework, brain activity is translated (decoded) into internal representations of a pre-trained deep neural network (DNN). Functional magnetic resonance imaging (fMRI) responses measured while a human subject viewed 1200 natural images were used as training data. Linear regression models were trained to predict unit activations of individual DNN units in the DNN that responds to the same images. The pre-trained VGG and CLIP models were used as the DNNs in this study. **(b)** Image reconstruction through Bayesian estimation. The seen or imagined image is reconstructed from the decoded DNN unit activations. The decoded DNN unit activations are incorporated into the Bayesian posterior distribution of the image via likelihood functions. A prior distribution constructed with a pre-trained neural network-based image generator model is used in the Bayesian estimation. An image sampled from the posterior distribution is considered a reconstruction.

In Bayesian estimation, sampling from a posterior distribution is often intractable; thus, its application in neural decoding has been limited. Although a few previous studies have introduced Bayesian estimation into letter^5^ and face^37^ image reconstruction, these were cases where the posterior distribution could be analytically obtained. Their approach cannot be straightforwardly applicable to cases with natural images. As an alternative approach, we used the stochastic gradient Langevin dynamics (SGLD) algorithm^38^ to sample images from the posterior distribution. Our results demonstrated that seen and imagined images were successfully reconstructed from brain activity, supporting the effectiveness of the SGLD algorithm in the field of neural decoding.

Subsequently, a pre-trained contrastive language–image pre-training (CLIP) model^39^ was used to leverage semantic information from the brain for image reconstruction. Because the CLIP model has been trained to obtain embeddings shared between images and their text captions in the last layer, the image encoder of the CLIP model is thought to extract semantic information from input images. As terminology, features and representations provided by the last layer of the CLIP model are called “semantic features” and “semantic representations,” respectively. Similarly, those provided by low/high layers of VGG19 are called “low/high-level visual features” and “low/high-level visual representations.” Here, using the same neural decoding procedure as that for VGG19, brain signals were translated into the semantic features provided by the CLIP model (Fig. 1a). Afterwhich, the decoded semantic features were introduced into our Bayesian image reconstruction framework through an additional likelihood function (Fig. 1b).

By applying the proposed framework to the dataset from Shen et al. (2019)^23^, we demonstrate that our framework can reconstruct seen images only using high-level visual information and externalize mental imagery.

## Results

### Reconstruction algorithm

The reconstruction framework presented in this study is based on the method described by Shen et al. (2019)^23^. In the previous study, fMRI signals from voxels in the visual cortex, measured while human subjects viewed images, were translated (decoded) into the hierarchical representations of VGG19 for the same images (Fig. 1a). For decoder construction, linear regression models were trained to predict the unit activations of individual DNN units in each layer of VGG19 using fMRI responses to 1200 natural images. The trained models (decoders) were then applied to the independent test data which consisted of fMRI signals measured while the same subjects viewed 50 natural images and 40 artificial shapes; and while, they imagined 10 natural images and 15 artificial shapes. For a given test trial, we denote the decoded representation (i.e., the decoded feature vector) of the *ι*-th VGG layer by 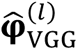. Using the decoded VGG representations from *L* layers, the observed or imagined image in the given test trial was reconstructed by solving the following optimization problem:

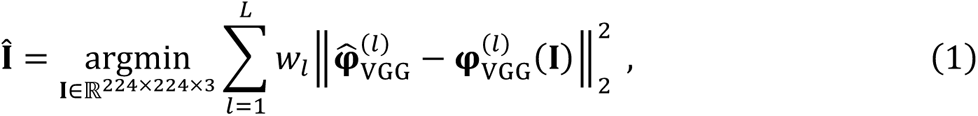

where 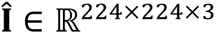 is the reconstructed image, *W_ι_* is a parameter determining the contribution of the *ι*-th VGG layer, and 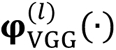 is the feature extraction function mapping the input image into the representation of VGG19’s ι-th layer. This optimization problem was solved using the momentum gradient method with some constraints. Whereas the previous study focused on the effect of combining multiple VGG layers, we attempted reconstruction with individual or a subset of layers in this study because the representations in high VGG layers could be more accurately decoded from the brain during imagery than low VGG layers. Thus, the VGG layers used in equation (1) were changed in our experiments.

In our proposed framework, the image is reconstructed in a Bayesian manner from 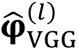. As the likelihood function, we use

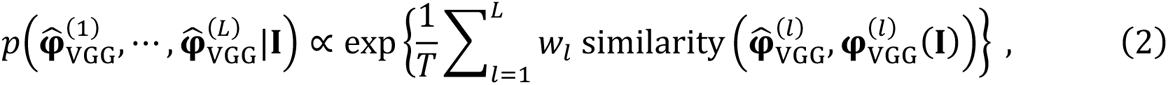

where *T* is a parameter called “temperature” and the function similarity(·,·) is a similarity metric to measure the similarity between two vectors; in this study, the Pearson correlation coefficient was used as the similarity metric. When the negative L2 norm is adopted as the similarity metric, the maximum likelihood estimation using this likelihood function is equivalent to the reconstruction method proposed by Shen et al. (2019). Thus, our proposed image reconstruction method is a Bayesian extension of Shen et al.’s (2019).

We used a pre-trained image generator model to prepare a prior distribution for the Bayesian estimation. Specifically, VQGAN^18^ trained on ImageNet training images^40^ was used in this study; however, other image generator models can be used in our framework. Given its latent vector **z**, the image generator model of VQGAN produces an image. We denote the probability distribution of the generated images conditioned on **z** by 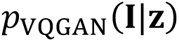. Note that the produced image **I** is deterministic with respect to **z** when we use VQGAN; however, here, we explain our framework with a probability distribution because our framework can also be combined with image generator models that probabilistically generate images. We can obtain an image prior *ρ*(**I**) by preparing a distribution *ρ*(**z**), constructing the joint distribution 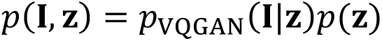, and marginalizing out **z**:

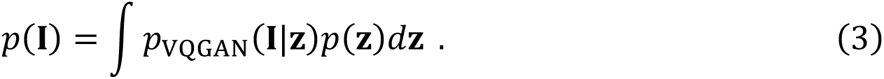

We used the non-informative distribution as *ρ*(**z**) in this study.

To achieve better reconstruction, we introduced the assistance of semantic information into our framework. Using the same decoding procedure as that for VGG19, given fMRI signals were translated into the unit activations in the last layer of the image encoder of the pre-trained CLIP model^39^. We denote this decoded unit activation vector by 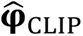. Using a likelihood function 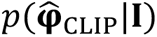 with a form similar to equation (2) (see Methods), the observed or imagined image was reconstructed from 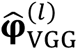 and 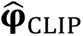 in a Bayesian manner (Fig. 1b). Thus, our posterior distribution was given by

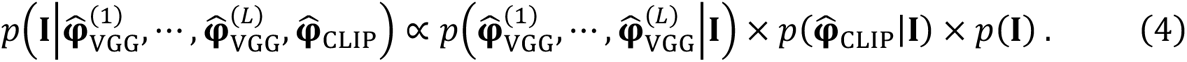

To obtain samples from the posterior distribution, we used the SGLD algorithm (see Methods). We performed 500 iterations and treated the obtained sample as the reconstructed image.

### Reconstruction of seen images

To assess the effectiveness of the proposed framework, we first applied it to the fMRI signals measured while three human subjects viewed 50 natural images and 40 artificial shapes. The data from Shen et al. (2019)^23^ were used. The decoders were trained using independent training data comprising fMRI signals measured while the same subjects viewed other 1200 natural images. Images reconstructed from the brain via the set of all VGG layers, via the set of conv2–fc6, and via the individual VGG layers are shown in Figs. 2a, 2b; Supplementary Figs. 1–3. These images were compared to those reconstructed using the method described by Shen et al. (2019). Whereas both methods produced moderately nice reconstructions using the full set of VGG layers, Shen et al.’s (2019) method failed to produce meaningful images without conv1, implying its reliance on low-level visual information; in contrast, meaningful images were obtained using the proposed framework.

**Fig. 2.**
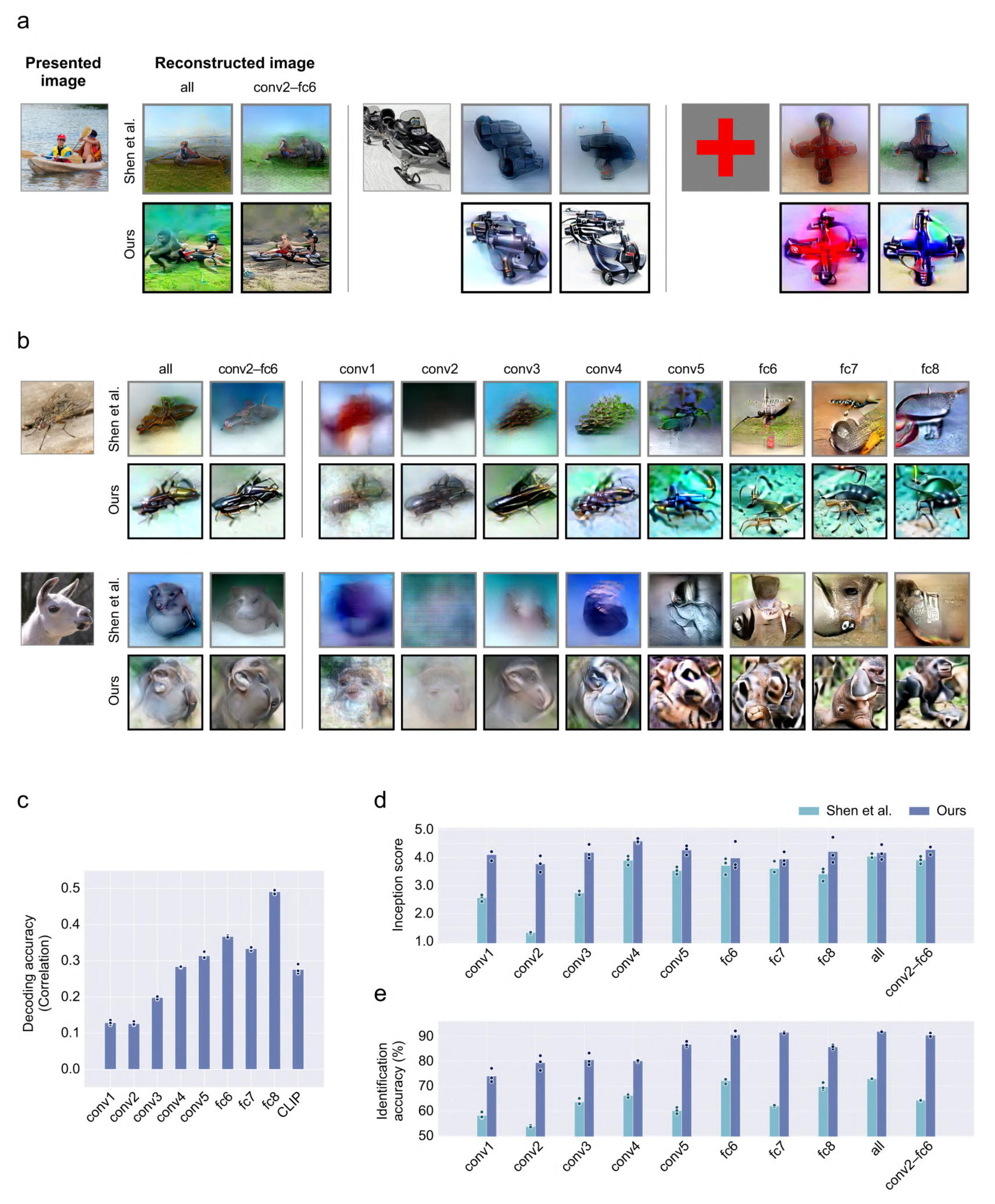
Reconstruction of seen images. **(a,b)** Reconstructed images. Seen stimulus images were reconstructed from brain-decoded unit activations. Images reconstructed through the set of all VGG layers, through the set of conv2–fc6, and through the individual VGG layers are shown here. The gray and black surrounding frames indicate images reconstructed using the method of Shen et al. (2019) and our proposed framework, respectively. All reconstructed images shown here are from Subject 2. Reconstructed images from individual subjects are provided in Supplementary Figs. 1–3. **(c)** Decoding accuracy. The decoding accuracy for each DNN unit was evaluated by computing the correlation coefficient between true and decoded unit activation values across 50 natural images. The mean accuracy across the DNN units in each layer is shown here. Black dots and blur bars indicate the mean accuracies for individual subjects and those across the three subjects, respectively. **(d)** Inception score. Light and dark blue bars indicate the mean Inception scores across the three subjects for images reconstructed using the method of Shen et al. (2019) and our proposed framework, respectively. Dots indicate the Inception scores for individual subjects. **(e)** Image identification accuracy. The formats are the same as those in (d).

Note that successful reconstructions were obtained even for artificial shapes, although the brain decoders were trained using only the brain responses to natural images (Fig. 2a, right; Supplementary Figs. 4–6). These results demonstrate that our reconstruction framework has a strong generalization ability for images in a new unknown domain, excluding the possibility that it generates images by virtually picking them from limited exemplars.

We also evaluated how accurately the unit activations in individual layers were decoded from the brain. We computed the correlation coefficient between the true and decoded unit activations across the 50 natural images for each unit, and then computed the mean correlation coefficient across the units in each layer (Fig. 2c). As in previous studies^23, 33^, all individual layers were decoded with moderate accuracy. The results suggest that the decoded unit activations of high VGG layers as well as low VGG layers carry a significant amount of information on the presented images.

To quantitatively evaluate the quality of the reconstructed images, we performed two types of evaluations. First, we computed the Inception score^41^ for these images to evaluate their visual quality (Fig. 2d). Our reconstructions demonstrated higher Inception scores than those of Shen et al. (2019). Second, to examine whether the reconstructed images preserved information about the seen images, we performed a pairwise image identification analysis. Following the procedures in previous studies ^3, 23^, we examined whether each reconstructed image was more similar to the corresponding seen image than a randomly selected one; and reported the proportions of correct answers (Fig. 2e). The weighted similarity in each reconstruction algorithm was used as the similarity metric (i.e., the sum in equation (2)). The identification accuracies of the proposed framework were higher than those of Shen et al. (2019). A similar tendency was consistently observed with the reconstructed images of the artificial shapes (Supplementary Fig. 7).

### Reconstruction of imagined images

We applied the reconstruction methods to the fMRI signals measured during imagery (Figs. 3a, 3b; Supplementary Figs. 8 and 9). We used the data measured while the subjects imagined 10 natural images and 15 artificial shapes. Although the reconstruction quality varied significantly across samples, our reconstruction framework successfully produced interpretable images that reflected the target images to be imagined; in contrast with Shen et al.’s (2019) method. A comparison of the decoding accuracy between the DNN layers demonstrated that the decoding accuracies for conv1 and conv2 were considerably low compared to those of the other layers (Fig. 3c), which is consistent with the view that our proposed framework leverages high-level visual information better than that proposed by Shen et al. (2019).

**Fig. 3.**
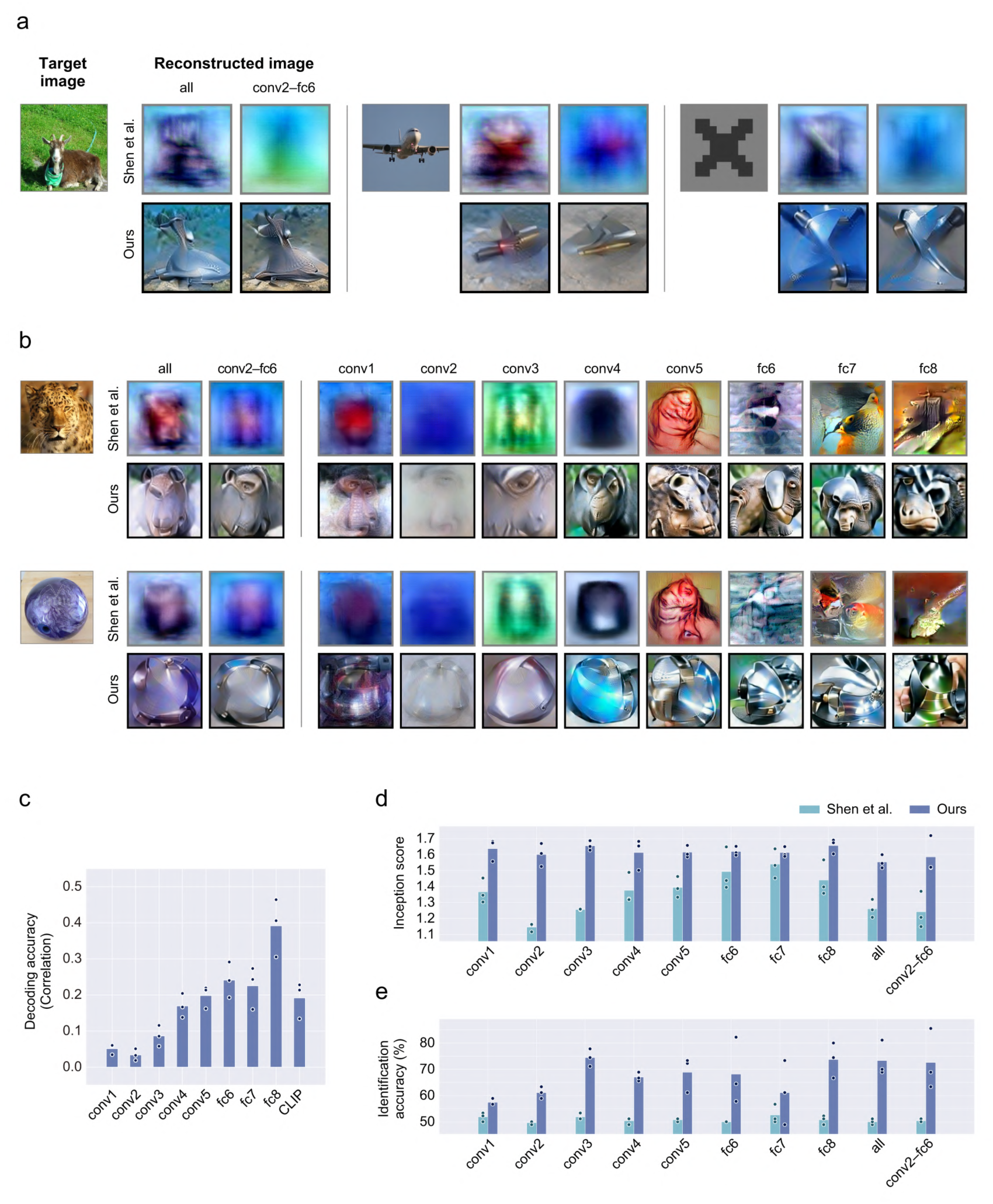
Reconstruction of imagined images. Imagined images were reconstructed from brain activity. The formats are the same as those in Fig. 2. All reconstructed images in panels (a) and (b) are from Subject 2. Those from the other subjects are provided in Supplementary Fig 8.

We evaluated the quality of the reconstructed images using the same procedure as that used for seen image reconstruction. Our proposed framework outperformed Shen et al.’s (2019) in terms of both Inception score and identification accuracy (Figs. 3d, 3e). Furthermore, our framework successfully reconstructed artificial shapes, although the brain decoders were trained using only brain responses to natural images (Supplementary Figs. 9 and 10), indicating its strong generalization ability.

Interestingly, we found that, for some artificial shapes, line components in imagery reconstructions were emphasized compared to those in seen image reconstructions. A comparison of the reconstructed images for an X-shaped geometric pattern between the imagery and the seen image reconstructions is shown in Fig. 4a. Lines with orientations of approximately 45° and 135° appear in the imagery reconstructions, while rough silhouettes of the target shape were emphasized in the seen image reconstructions. This tendency was consistently observed across the three subjects and five different colors (Supplementary Fig. 11). To quantitatively assess this tendency, we quantified how strongly the line components with each orientation were included in the reconstructed images. Briefly, following the procedure described in previous studies^28, 42^, the strengths of the line components for individual orientations in a reconstructed image were evaluated by applying the Radon transform to the image (see Methods). The imagery reconstructions had stronger line components at orientations of 45° and 135° than the seen image reconstructions (Fig. 4b). These results may reflect the sharpening effect caused by the top-down process in the brain^43^. We conducted a supplementary analysis to investigate the contribution of each brain subarea in the visual cortex to the reconstruction. Following previous studies^23, 44^, we divided the visual cortex into five subareas: V1, V2, V3, V4, and the higher visual cortex (HVC). We then performed the same decoding and reconstruction procedures while limiting the input brain area to each of those five subareas (Supplementary Fig. 12). HVC outperformed the other subareas, indicating that HVC made the largest contribution to imagery reconstruction.

**Fig. 4.**
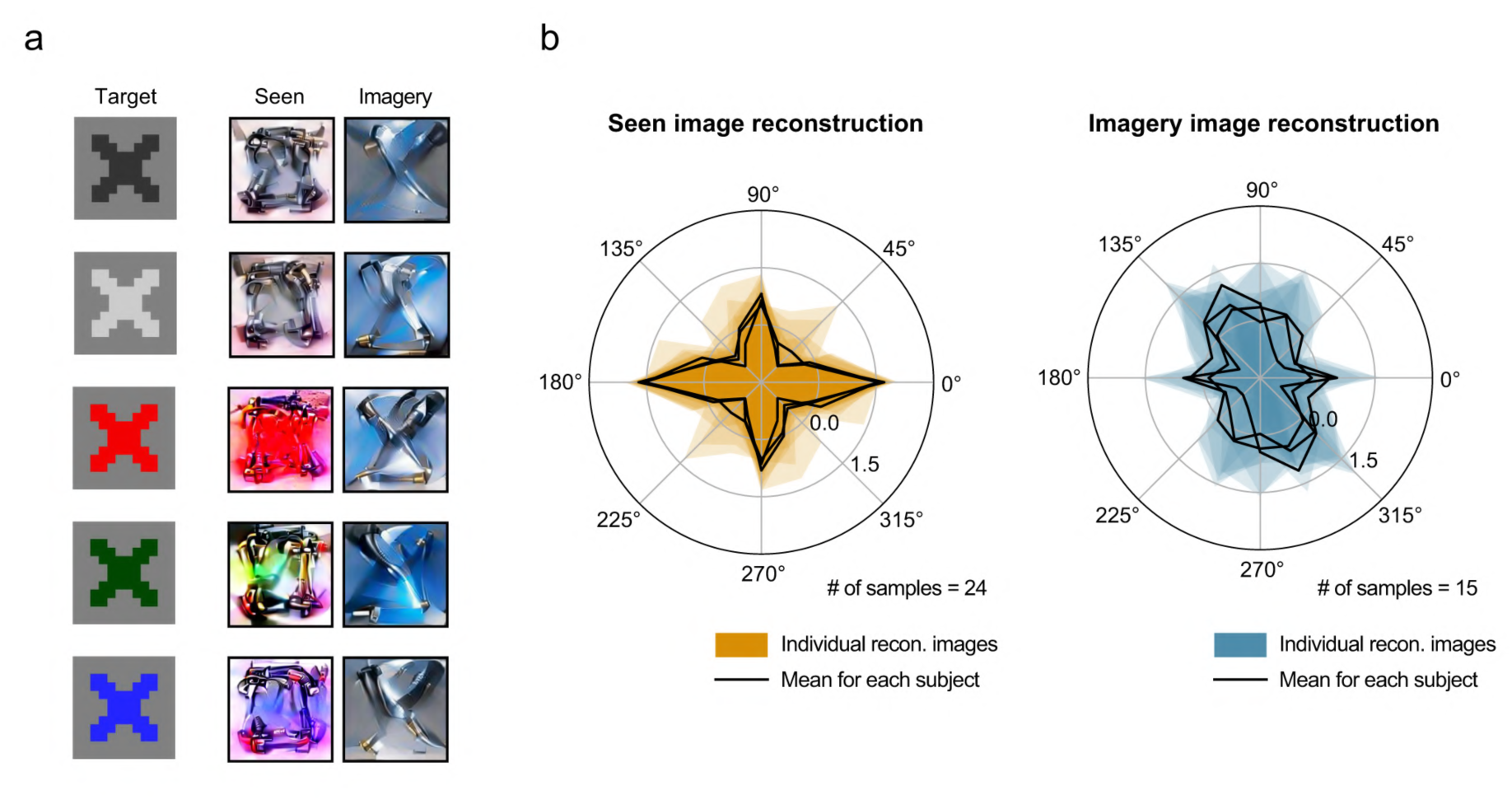
Comparison of seen image reconstructions and imagery reconstructions. **(a)** Reconstructions of an X-shaped geometric pattern. Images reconstructed from brain activity measured while Subject 2 viewed X-shapes and images reconstructed from brain activity measured while the same subject imagined the same X-shapes are compared. All reconstructed images shown here are from Subject 2. Those from the other subjects are provided in Supplementary Fig. 11. **(b)** Quantitative evaluation. The strength of line components with each orientation in individual reconstructed images for X-shapes was evaluated as a quantitative assessment. The results for seen image reconstructions (left) and those for imagery reconstructions (right) are shown in polar plots.

### Effect of the image prior

To characterize the effect of the image prior, we performed an ablation analysis. To reconstruct images without the image prior, we conducted maximum likelihood estimation using the likelihood function in equation (4) (Figs. 5a, 5b). The fMRI data measured while the subjects imagined natural images were used for this analysis. The appearances of the reconstructed images using our full method were significantly better than those obtained without the image prior. The quantitative comparison of the Inception score and image identification accuracy also supported this tendency (Figs. 5c, 5d). We also observed the same tendency with the seen image reconstructions (Supplementary Fig. 13)

**Fig. 5.**
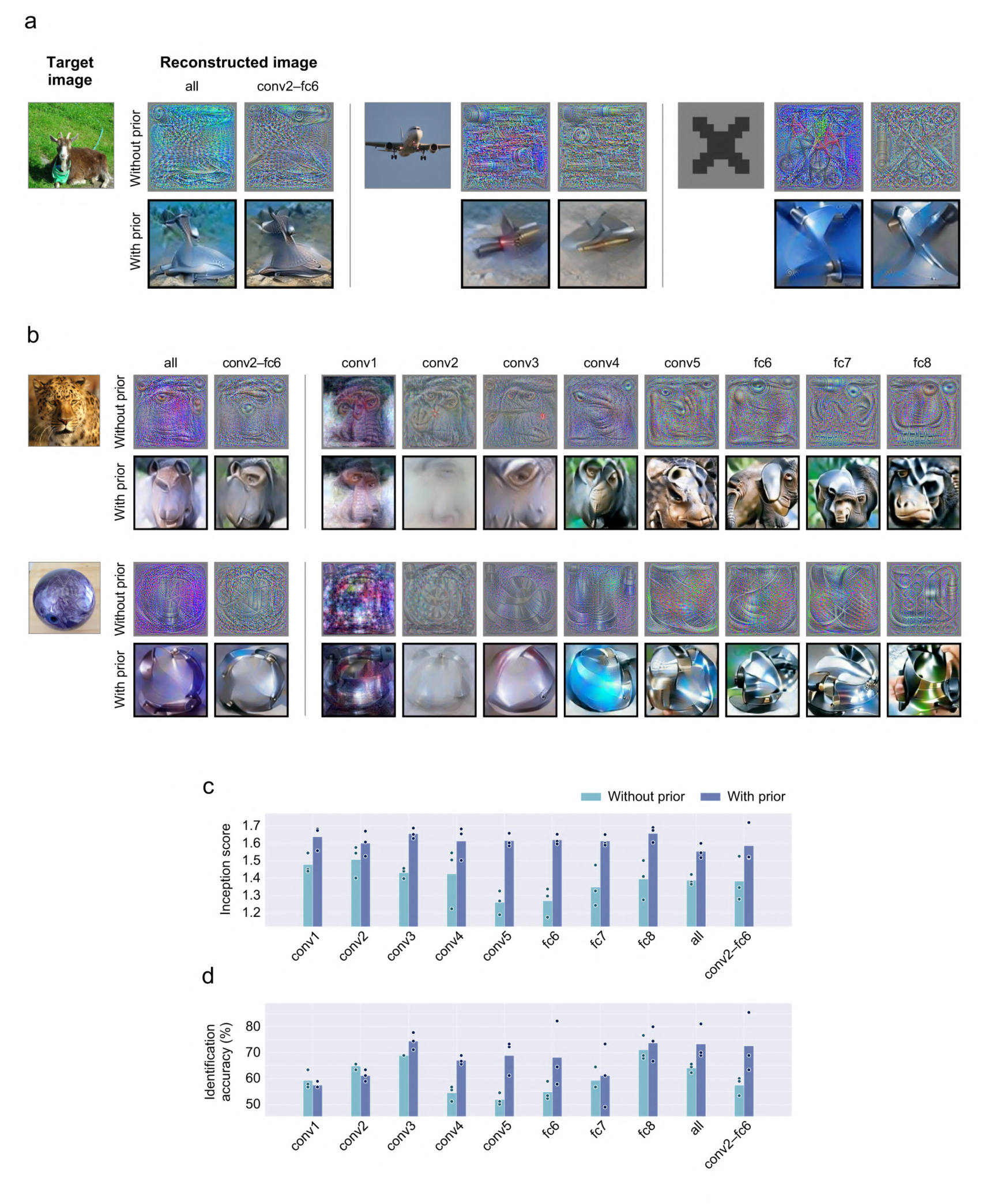
Effect of the image prior. Imagery reconstruction results are compared between our full method and that without the image prior. The formats and target images are the same as those in Fig. 3.

### Effect of the assistance of semantic information

We performed an ablation analysis to characterize the effects of the semantic information assistance. Here, we gradually varied the value of the hyperparameter controlling the strength of the influence of the CLIP features (coefficient λ_CLIP_; see Methods for details), and reconstructed the images. λ_CLIP_ = 0 indicates no assistance; whereas a higher coefficient introduces stronger assistance into the reconstructed images. λ_CLIP_ = 0.25 was used as the default. The imagery reconstructions were compared across different coefficient values (Fig. 6a). Removing the assistance of semantic information resulted in a drop in the image identification accuracy while the Inception score was almost maintained (Figs. 6b, 6c). Interestingly, small or no improvements were observed for seen image reconstructions (Supplementary Fig. 14), suggesting that this semantic assistance compensates for the lack of low-level visual information in the imagery reconstruction.

**Fig. 6.**
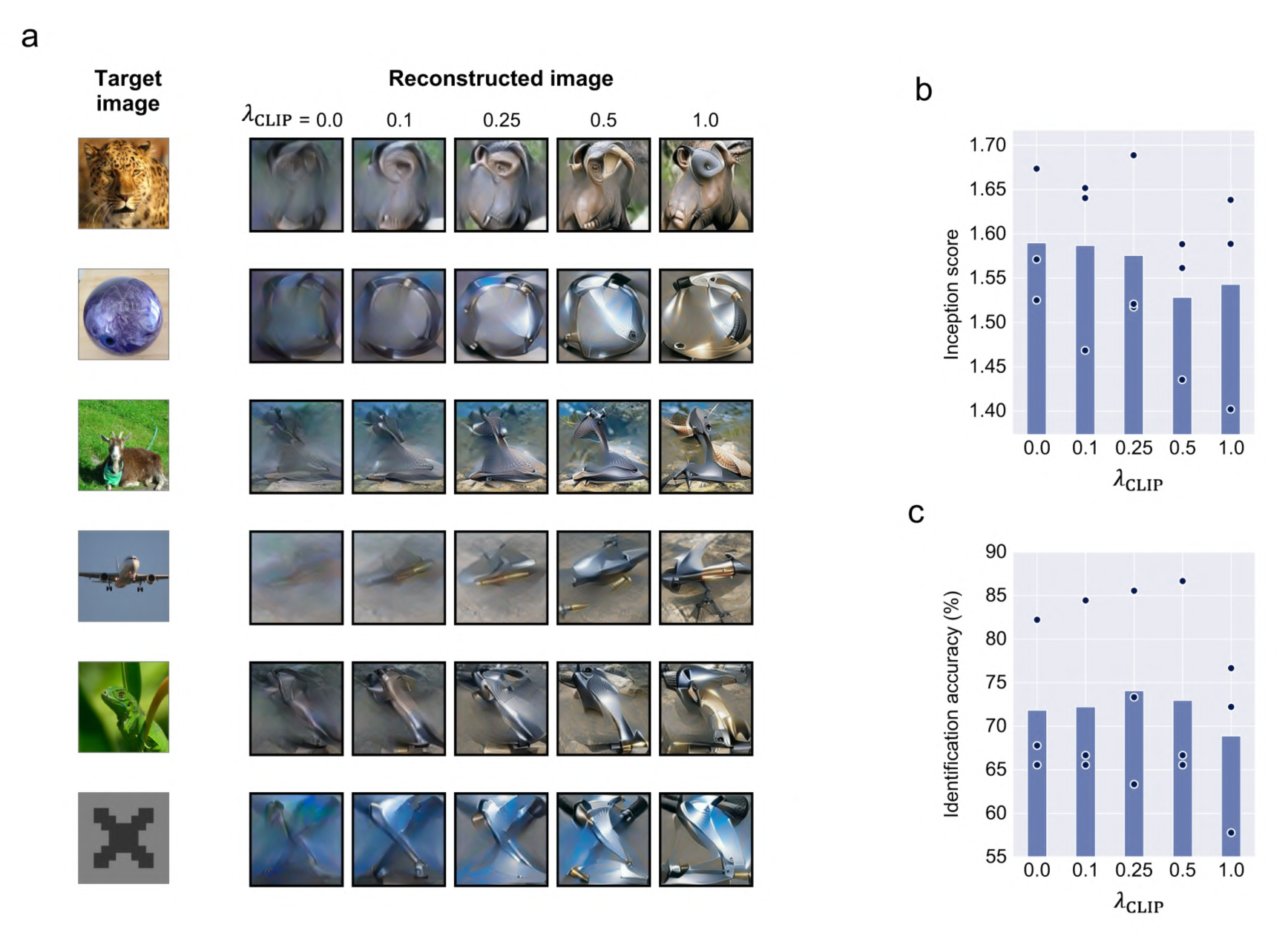
Effect of the assistance of semantic information. **(a)** Imagery reconstructions with different strengths of semantic information assistance. The hyperparameter controlling the strength of semantic information assistance varied from 0.0 to 1.0. Images reconstructed via conv2–fc6 are shown. **(b)** Inception score. **(c)** Image identification accuracy.

## Discussion

To reconstruct mental imagery from brain activity, we extended the previous image reconstruction method proposed by Shen et al. (2019)^23^ to a Bayesian estimation framework and introduced the assistance of semantic information (Fig. 1). While this previous reconstruction method was highly dependent on low-level visual information (i.e., low-layer VGG representations) decoded from the brain, our current proposed framework successfully produced meaningful reconstructions only using high-level visual information (Fig. 2). This advantage allows for the better use of such high-level visual information retained in the brain during imagery, thereby enabling mental image reconstruction (Fig. 3). Subsequent ablation analyses demonstrated that the two components introduced into our framework (Bayesian estimation and assistance of semantic information) were necessary for meaningful reconstructions (Figs. 5 and 6).

Our framework could reconstruct artificial shapes, even though the brain decoders were solely trained with fMRI responses to 1200 natural images (Figs. 2 and 3, Supplementary Figs. 4–7, 9, and 10). These results demonstrate that our reconstruction framework has a strong generalization ability for images in a new unknown domain, excluding the possibility of generating images by virtually selecting them from limited exemplars. As our spontaneous thoughts are not controlled or limited in daily life, such a strong generalization ability is potentially helpful for brain–machine interface applications in practical situations. In addition, the Bayesian nature of our framework would allow for better reconstruction by using a customized image prior when the domain of the image to be reconstructed is known in advance. According to a previous study^5^, using a suitable prior improves the reconstruction of hand-written letter images from brain activity. Exploring this Bayesian advantage with a variety of image domains or sensory modalities other than vision would be an important challenge in the field of neural decoding.

We used the stochastic gradient Langevin dynamics (SGLD) algorithm^38^ to obtain samples (i.e., reconstructed images) from the Bayesian posterior distribution. In Bayesian estimation, sampling from the posterior distribution is often intractable. Thus, the use of Bayesian estimation for visual image reconstruction has been limited. A few previous studies have introduced Bayesian estimation into visual image reconstruction, which can be divided into two types: 1) cases in which the posterior distribution is analytically obtained, and 2) cases in which the maximum a posteriori probability estimate is approximately obtained by selecting it from a finite set of samples. For example, Schoenmakers et al. (2013)^5^ adopted the first approach and demonstrated that their Bayesian framework produced better reconstructions of hand-written letter images. In this case, a multivariate Gaussian distribution was used as the image prior. A similar concept has been applied to face image reconstruction in a subsequent study^37^. However, given that the posterior distribution must be obtained analytically, this first approach cannot be combined with more general and flexible priors. In the second approach, a set of images is independently prepared in advance, and the image with the highest posterior probability is treated as the reconstructed image^10, 45^. Theoretically, this method would work if an infinite or sufficient number of samples have been prepared, but cannot reconstruct arbitrary images with limited exemplars. In our study, we introduced the SGLD algorithm into the reconstruction framework as an alternative approach and demonstrated that an image prior constructed with a pre-trained neural network improved the quality of reconstructions (Fig. 5). These results demonstrate the effectiveness of the SGLD algorithm for neural decoding.

The image generation process of our framework resembles that of a text-to-image generation method. In a popular algorithm for text-to-image generation, an image is generated by optimizing the latent vector of a pre-trained image generator model such that the output image matches the target text in the multi-modal embedding space provided by the CLIP model^46^. Thus, although our reconstruction framework was derived from Shen et al. (2019), it can be considered an extension of such a text-to-image generation algorithm for brain-to-image generation. Additionally, brain-to-text generation as a fusion of the above two types of algorithms would be an interesting future topic in the field of neural decoding.

While our reconstruction framework provides fundamental technology for brain–machine interfaces, it also serves as a tool for investigating the generation process of mental imagery. The comparison of input brain areas in the human visual hierarchy showed that the highest quality of imagery reconstruction was achieved with the higher visual cortex (HVC) (Supplementary Fig. 12). These results are consistent with previous neuroimaging studies supporting the idea that HVC is recruited more than the lower visual areas during imagery. Furthermore, we found that the line components in the imagery reconstructions of some artificial shapes were emphasized compared to those in the seen image reconstructions (Fig. 4). Although further investigation is required, this finding probably reflects the sharpening effect caused by the top-down process in the brain^43^. Therefore, our framework provides a novel approach for investigating hypotheses regarding mental imagery.

## Methods

### fMRI dataset

We used the fMRI dataset from a previous study^23^, which can be downloaded from the Figshare repository (https://figshare.com/articles/dataset/Deep_Image_Reconstruction/7033577). The dataset comprises fMRI data from three human subjects (Subjects 1–3). In this experiment, each subject viewed or imagined an image in each trial, and the brain activity was measured using fMRI. The fMRI data were divided into two sets: training and test datasets. The training dataset was used for decoder training and the test dataset was used for evaluation in the previous study. The same data splitting approach was adopted in the present study. The training dataset comprises fMRI data measured while the subjects viewed 1200 natural images. Each image was presented to each subject five times. Thus, 6000 fMRI responses per subject were available as training data. The test dataset comprises fMRI data measured while the subjects viewed 50 natural images and 40 artificial shapes (geometric shapes) and those measured while the subjects imagined 10 natural images and 15 artificial shapes. For this test dataset, each subject viewed each natural image 24 times and each artificial shape 20 times, and each subject imagined each natural image 20 times and each artificial shape 20 times. To adopt the same fMRI preprocessing procedure as in the previous study, we downloaded the preprocessed fMRI data from the Figshare repository. Following the same procedure, the training data were used without trial averaging and the trial-averaged test data were used for evaluation.

### Pre-trained neural networks

Three pre-trained neural networks were used in this study: VGG19^13^, VQGAN^18^, and CLIP’s image encoder^39^. We used a pre-trained VGG19 model provided by PyTorch. The outputs (unit activations) from conv1_2, conv2_2, conv3_4, conv4_4, conv5_4, fc6, fc7, and fc8 layers were used as the hierarchical representations in the brain decoding analysis. Following the procedure by Shen et al. (2019), the unit activation values before rectification were used as the targets to be decoded. In this study, these eight layers are called conv1, conv2, conv3, conv4, conv5, fc6, fc7, and fc8, and the unit activation vector of the *ι*-th layer for an input image 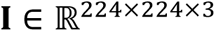 is denoted by 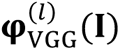.

A pre-trained VQGAN model was downloaded from the official GitHub repository (https://github.com/CompVis/taming-transformers). The model “VQGAN ImageNet (f=16), 1024” was used in this study. VQGAN uses a latent vector **z** as the input and produces an image as the output. The probability distribution of the output image given **z** is denoted by 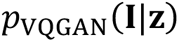. Note that the output of VQGAN is deterministic with respect to **z**. However, we describe our framework using a probability distribution because our framework can also be combined with image generator models that probabilistically produce images.

The CLIP image encoder was downloaded from the official GitHub repository (https://github.com/openai/CLIP). The model “ViT-B/32” was used in this study. The output from the last layer was used as the target to be decoded and denoted by 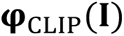.

Conventionally, the features and representations provided by low/high layers of VGG19 are called “low/high-level visual features” and “low/high-level visual representations”. Similarly, those provided by the last layer of CLIP are called “semantic features” and “semantic representations” in this study.

### Brain decoder

Brain activity measured using fMRI was translated (decoded) into hierarchical representations of VGG19 (Fig. 1a). For decoder construction, linear regression models were trained to predict the unit activations of individual units in each layer of VGG19 using the training dataset (i.e., fMRI responses to 1200 natural images). We used the linear regression algorithm with L2-regularization. Unless stated otherwise, fMRI signals from the voxels in the whole visual cortex were used as input for predicting the layers with spatial dimensions (i.e., conv1–conv5), because all individual subareas in the visual cortex are known to have considerable spatial information^47^. To predict the layers without spatial dimensions (i.e., fc6– fc8), fMRI signals from the voxels in the higher visual cortex (HVC) were used. According to previous studies, these layers can be accurately predicted from fMRI signals in HVC, and this choice is expected to reduce the risk of overfitting^33, 44^. Before linear regression training, fMRI voxels (i.e., input dimensions) were selected using the following voxel selection procedure. A customized, computationally expensive sparse algorithm was used in Shen et al. (2019), but we adopted the following procedure for computational efficiency.

Input voxel selection was performed using the training dataset to reduce the computational time and the risk of overfitting. To predict a given VGG layer, we applied principal component analysis to its representations across the 1200 images and extracted the principal components that explained more than 99% of the variance. Subsequently, for each voxel, we computed the correlation coefficients between the fMRI signal and the individual principal components. The maximum absolute value of the correlation coefficients was assigned to the voxel. This procedure was repeated for all voxels, and the voxels were ranked in descending order of the assigned correlation values. The top *k* voxels were used as inputs for the L2- regularized linear regression algorithm. The activation of each unit in the layer was predicted from the fMRI signals in the selected voxels. The number of voxels used (*k*) and the regularization parameter were optimized by cross-validation using the training dataset.

The trained linear regression models (brain decoders) were applied to fMRI data in the test dataset. The decoded representations for a given test trial are denoted by 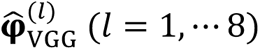 in this study. The same decoding procedure was also performed to predict the last layer of CLIP, and the decoded representation is denoted by 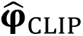.

In the analysis shown in Supplementary Fig. 12, to compare the reconstruction quality between subareas in the visual cortex, we performed the above decoding procedure using only fMRI signals in each of V1, V2, V3, V4, and HVC. We selected the voxels in each subarea using the labels provided with the preprocessed fMRI data from the Figshare repository.

### Proposed reconstruction framework

This section describes the exact form of the proposed reconstruction algorithm. The seen or imagined image was reconstructed from 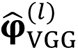 and 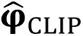 using a Bayesian estimation framework.

We assume that the log likelihood functions are given by

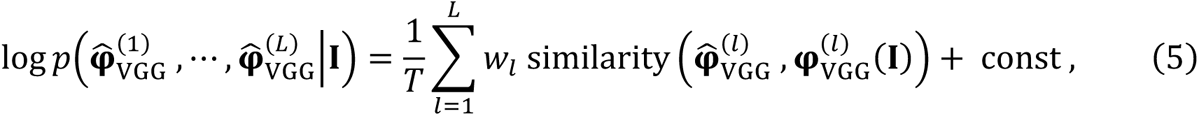

and

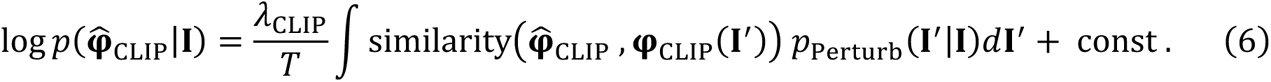

Here, *T* is a parameter called “temperature”, and the function similarity(·,·) is a similarity metric to measure the similarity between two input vectors. The Pearson correlation coefficient was used as the similarity metric in this study. *T* was set to 10^-7^. In equation (5), *W*_ι_ is a parameter controlling the strength of the contribution from the *ι*-th VGG layer, and this was set to the inverse of the number of used VGG layers. The reconstruction was performed with each VGG layer, with a subset of the VGG layers, or with all VGG layers. When a single layer or a subset of the layers was used, the sum in equation (5) was taken across the layer(s). For equation (6), λ_CLIP_ is a parameter controlling the strength of the semantic assistance. It was set to 0.1 and 0.25 for seen image reconstruction and imagery reconstruction, respectively. To make the likelihood function robust to perturbations in **I**, a random affine transformation is applied to **I**, and the mean similarity is evaluated in the likelihood function. The conditional probability distribution 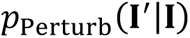 represents the random affine transformation, and we set its details according to the settings adopted in an implementation of text-to-image generation (https://medium.com/geekculture/text-to-image-synthesis-using-multimodal-vqgan-clip-architectures-896b8a6588ef). When we use the stochastic gradient Langevin dynamics (SGLD) algorithm, the mean in equation (6) is replaced by the empirical mean with 32 random samples.

An image prior consisting of a pre-trained image generator model was used in our framework. We used VQGAN as the image generator model. Optionally, other image generator models can be introduced into this framework. A pre-trained image generator model produces an image using a latent vector **z** as the input. We denote this relationship by 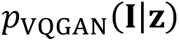. By preparing a distribution of **z**, the joint distribution of **I** and **z** can be defined. We denote it by 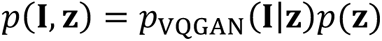, and denote the marginal distribution of **I** by ρ(**I**). Using ρ(**I**) as a prior distribution, we sample an image from the posterior distribution 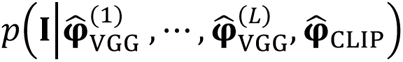 and the obtained image is treated as a reconstructed image. Any analytically differentiable distribution ρ(**z**) can be used in this framework. In this study, we assumed it to be the non-informative distribution.

To obtain an image from the posterior distribution, we used the SGLD algorithm^38^. The proposed posterior distribution can be rewritten as follows:

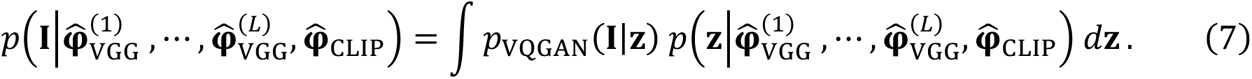

Thus, if we can sample **z** from 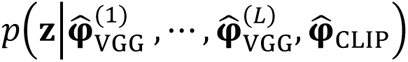, we can sample an image from the above distribution through 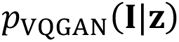. We performed sampling from 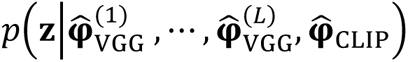 using the SGLD algorithm. The update rule of the algorithm is given by

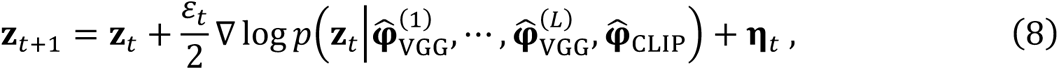

where **z**_t_ and Ɛ_t_ are the sample and the learning rate at step *t*, and ƞ_t_ is a vector whose elements are sampled from the Gaussian distribution *N*(0, Ɛ_t_). In this study, the learning rate was gradually decreased, and its sequence was given by

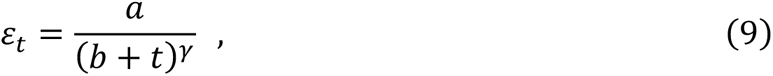

where *a*, *b*, and γ were set to 0.00015, 0.15, and 0.055. We performed 500 iterations, and a reconstructed image was produced using **z**_500_. For fast convergence, first we performed the Adam algorithm with an initial random latent vector. Then, the SGLD algorithm was started using the result from the Adam algorithm as the initial latent vector.

At every update in the SGLD algorithm, we approximately computed the right-side of equation (8). The logarithm of 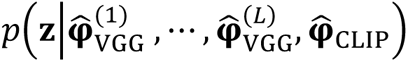 was represented by

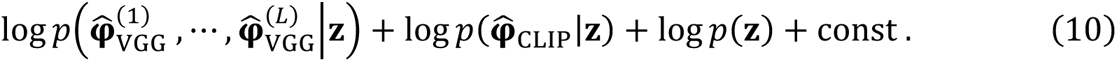

To approximately compute the first term, we rewrote the first term as

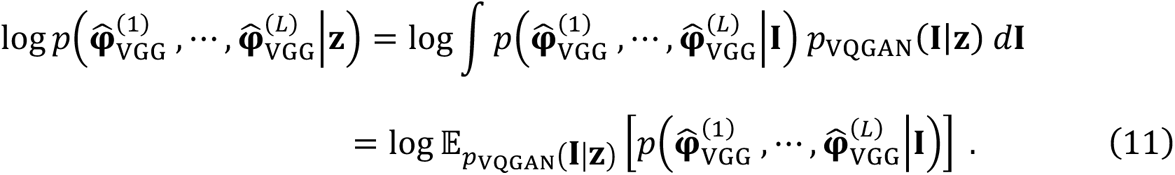

Using Jensen’s inequality, this term was approximated by

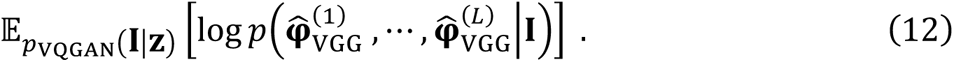

The inside of the square brackets is equivalent to equation (5); therefore, equation (12) was approximately computed by replacing the expectation with the empirical mean. The gradient of equation (12) could also be approximately computed using the back-propagation algorithm. The same approximation procedure could be applied to the second term in equation (10). With these approximated gradients, the update procedure shown in equation (8) was performed and the reconstructed image was obtained in our reconstruction framework.

### Reconstruction algorithm proposed by Shen et al. (2019)

The reconstructed images obtained using our framework were compared to those obtained using the method proposed by Shen et al. (2019)^23^. We downloaded a Python implementation of their method and their decoded features from the official GitHub and Figshare repositories (https://github.com/KamitaniLab/DeepImageReconstruction, https://figshare.com/articles/dataset/Deep_Image_Reconstruction/7033577). All reconstruction results obtained using the method of Shen et al. (2019) in this study were obtained using these resources.

### Quantification of the strength of line components in reconstructed images

The strengths of the line components for each orientation in reconstructed images were evaluated. We used the procedure described in previous studies^28, 42^. We converted a given reconstructed image into grayscale, subtracted the mean pixel value from the individual pixels, and squared the pixel values. Subsequently, the Radon transform was applied to the resulting image. The result is denoted by *R*(*r*, θ), and following the previous studies, the strength of line components with an orientation of θ was quantified by Var_r_[*R*(*r*, θ)]. These values are shown in Fig. 4b.

## Data availability

We used the fMRI data from Shen et al. (2019)^23^. These are available in an open data repository (https://figshare.com/articles/dataset/Deep_Image_Reconstruction/7033577).

## Code availability

The codes for our proposed reconstruction framework will be available at our GitHub repository (https://github.com/nkmjm/mental_img_recon) soon after the publication of the journal version of this article. All resources used to reproduce the results of Shen et al. (2019) are available from open repositories (https://github.com/KamitaniLab/DeepImageReconstruction, https://figshare.com/articles/dataset/Deep_Image_Reconstruction/7033577).

## Author contributions

NK and KM designed the study; NK and KM proposed the algorithms; NK performed data analyses; NK, SN, and KM interpreted the data; NK, SN, and KM wrote the manuscript.

## Acknowledgments

The authors express their gratitude to Susumu Saito, Hidehiko Takahashi, Shuntaro Aoki, Fan Cheng, Yukiyasu Kamitani, and members of Kamitani laboratory at Kyoto university.

## Supplementary information for

**Supplementary Fig. 1.**
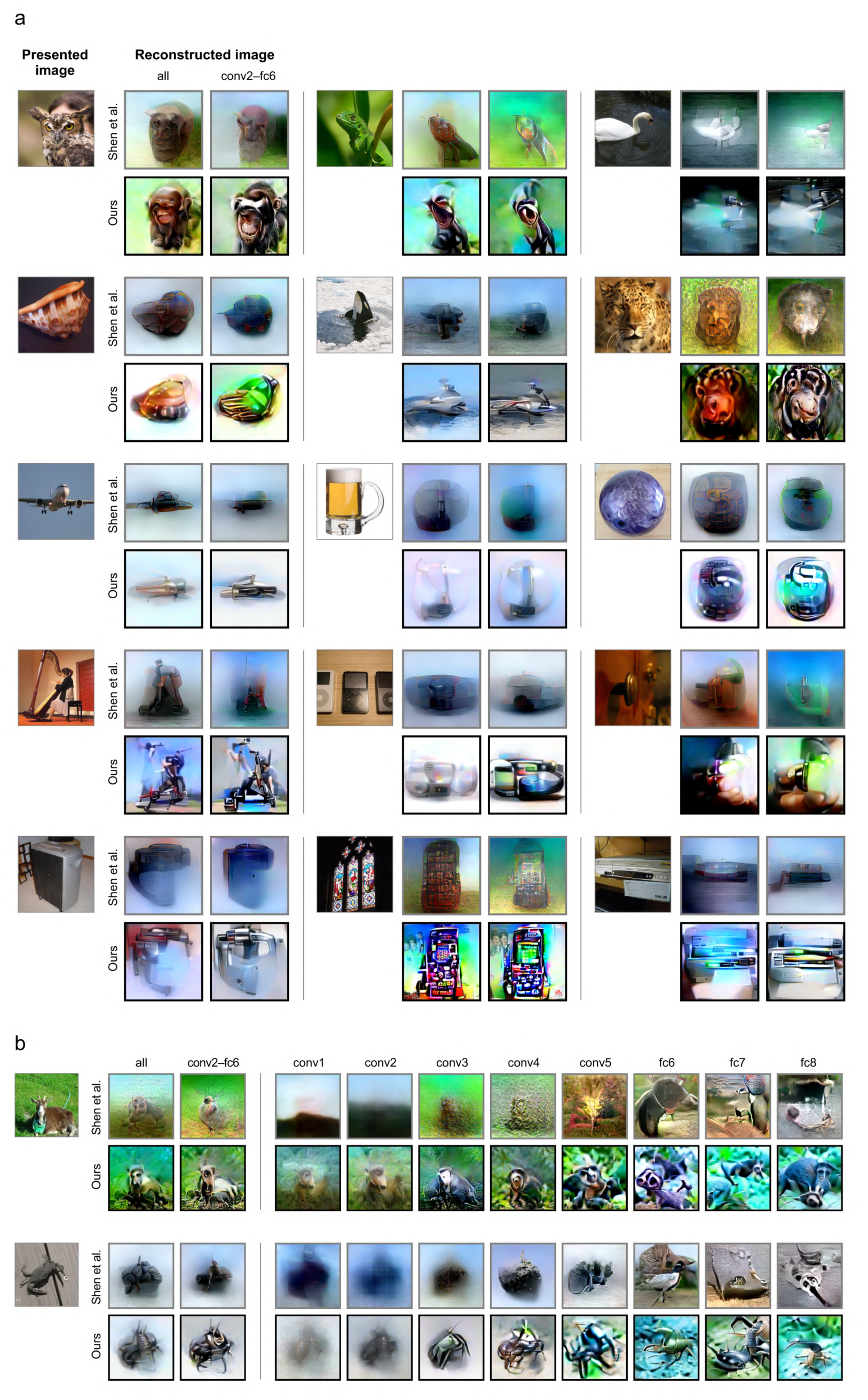
Reconstructions of seen images from Subject 1. The formats are the same as those in Figs. 2a and 2b.

**Supplementary Fig. 2.**
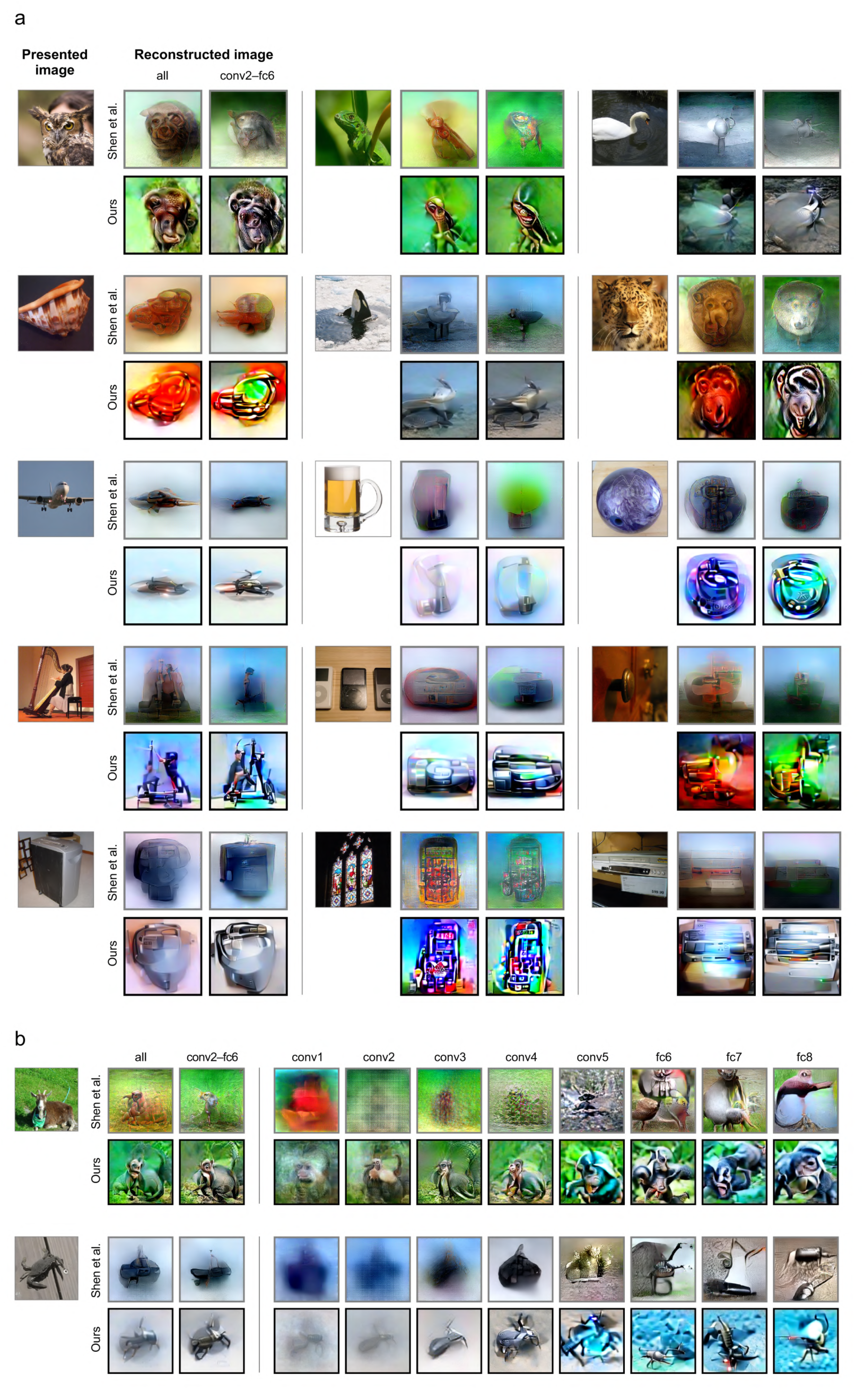
Reconstructions of seen images from Subject 2. The formats are the same as those in Figs. 2a and 2b.

**Supplementary Fig. 3.**
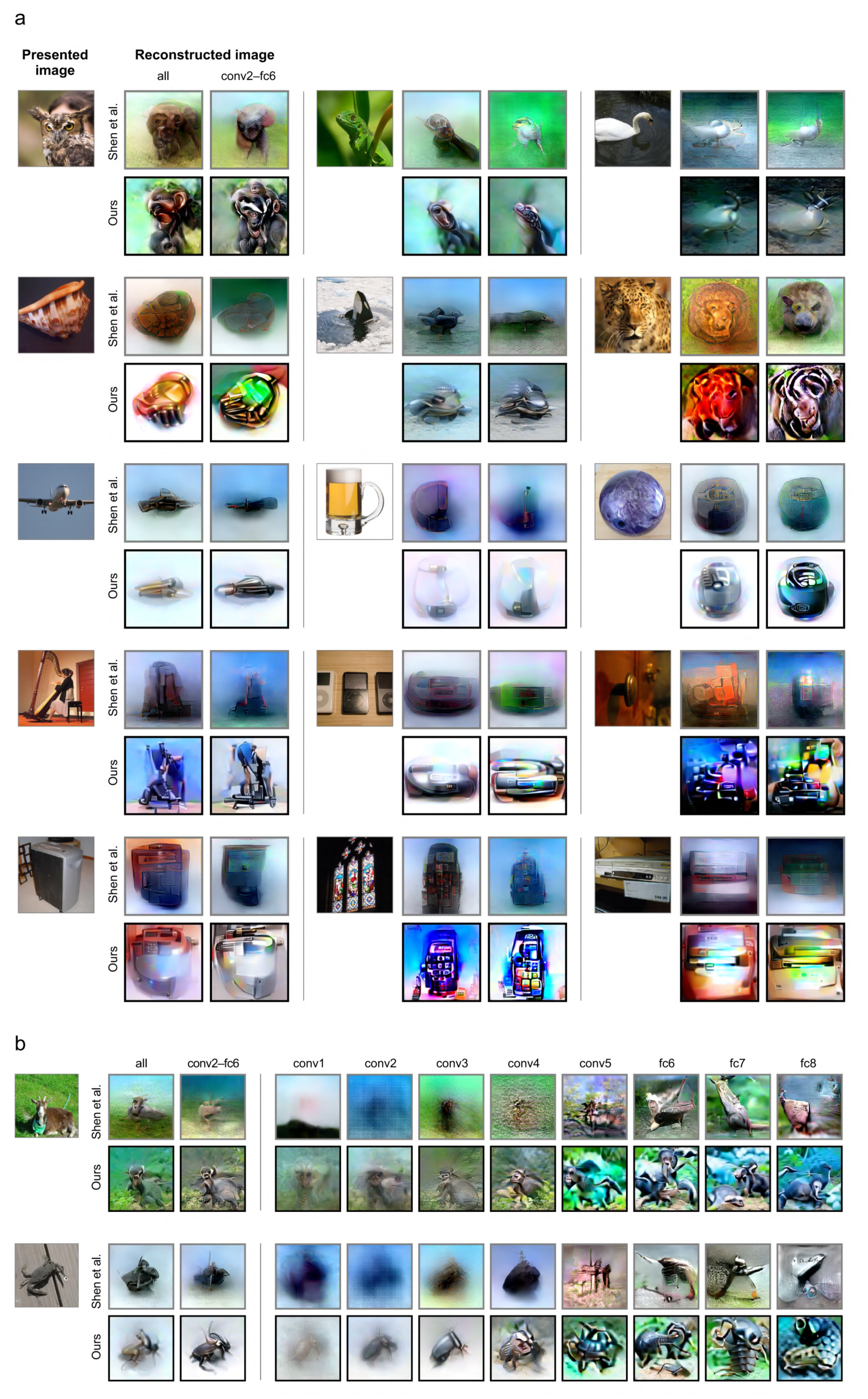
Reconstructions of seen images from Subject 3. The formats are the same as those in Figs. 2a and 2b.

**Supplementary Fig. 4.**
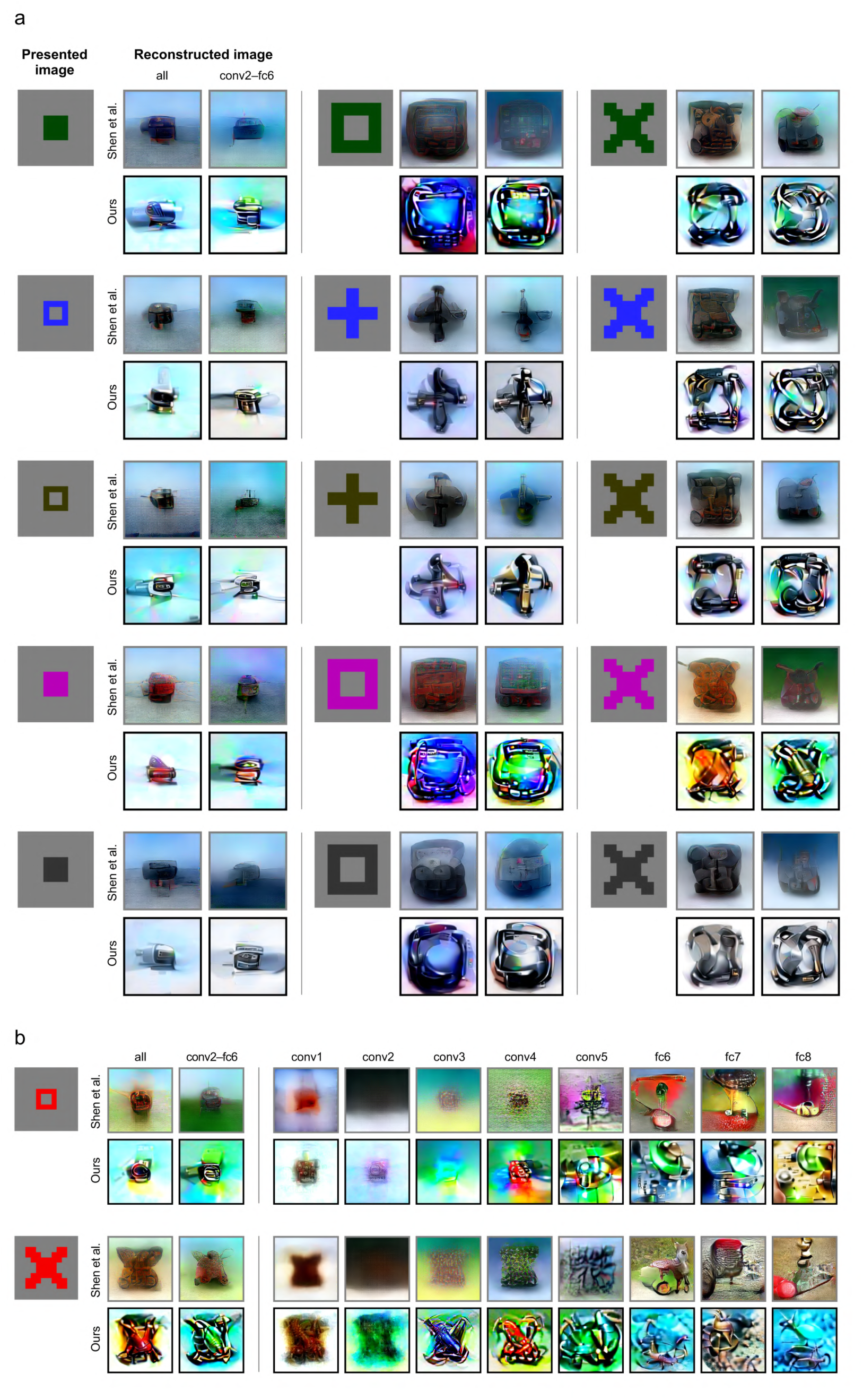
Reconstructions of seen artificial shapes from Subject 1. The formats are the same as those in Figs. 2a and 2b.

**Supplementary Fig. 5.**
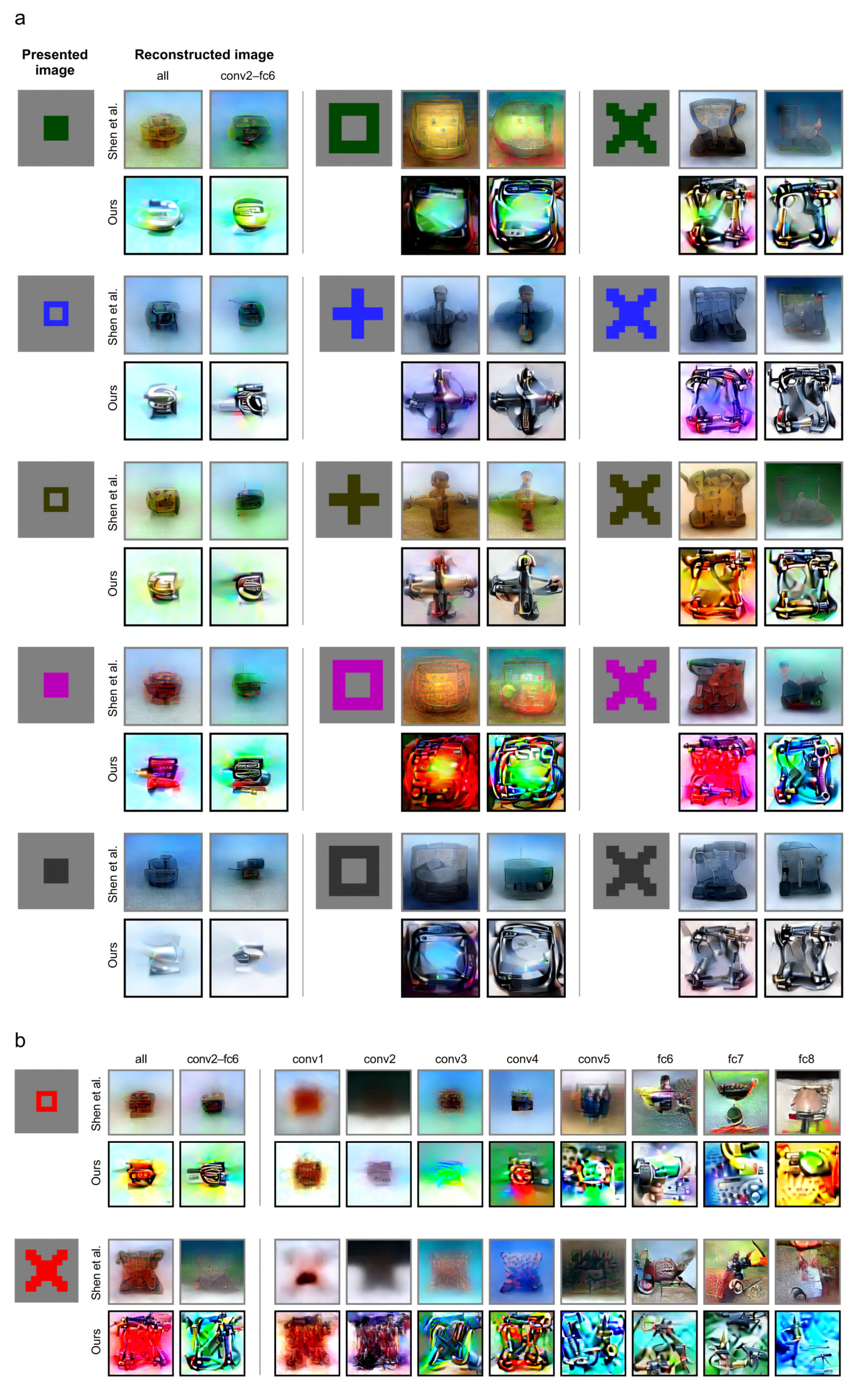
Reconstructions of seen artificial shapes from Subject 2. The formats are the same as those in Figs. 2a and 2b.

**Supplementary Fig. 6.**
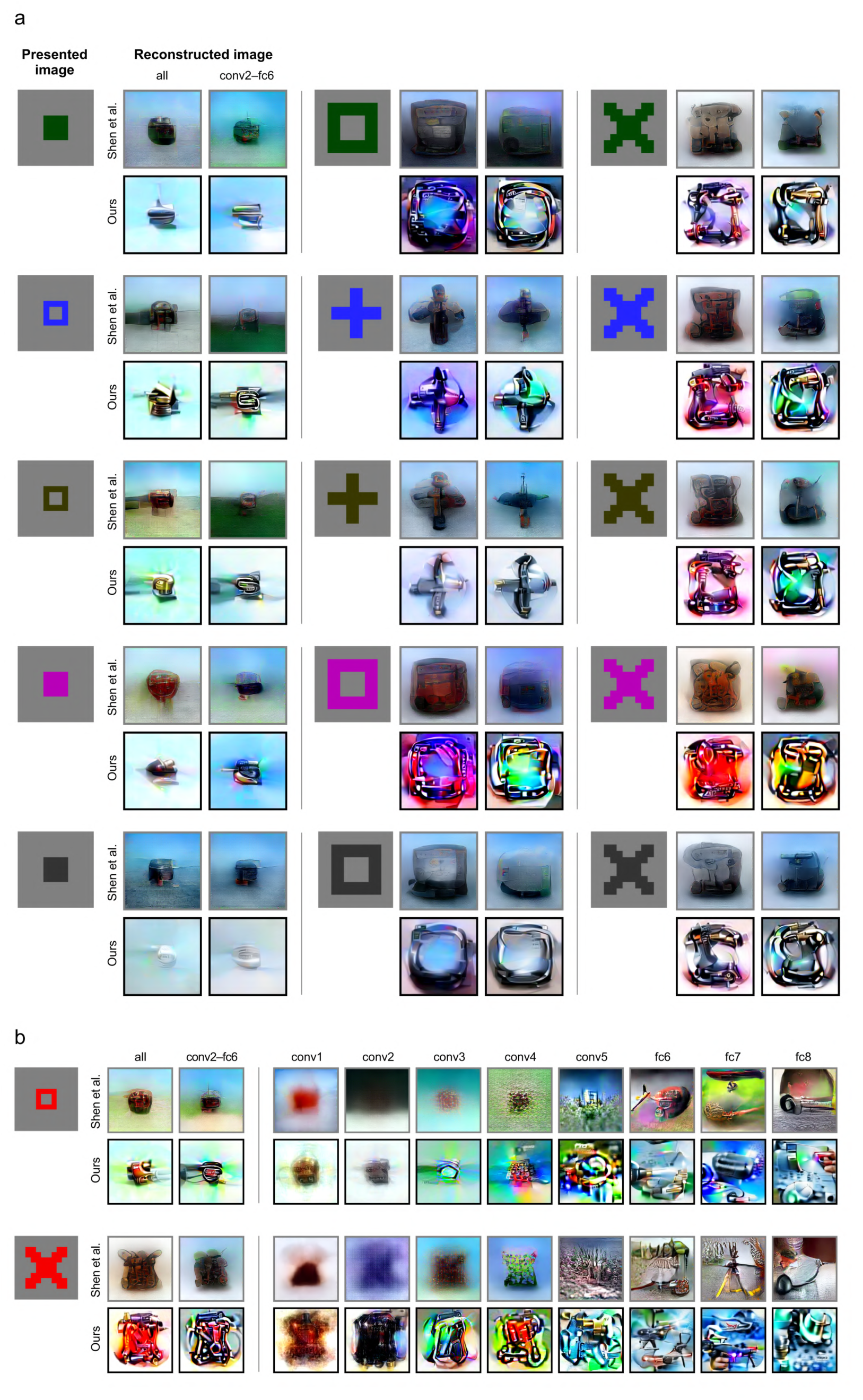
Reconstructions of seen artificial shapes from Subject 3. The formats are the same as those in Figs. 2a and 2b.

**Supplementary Fig. 7.**
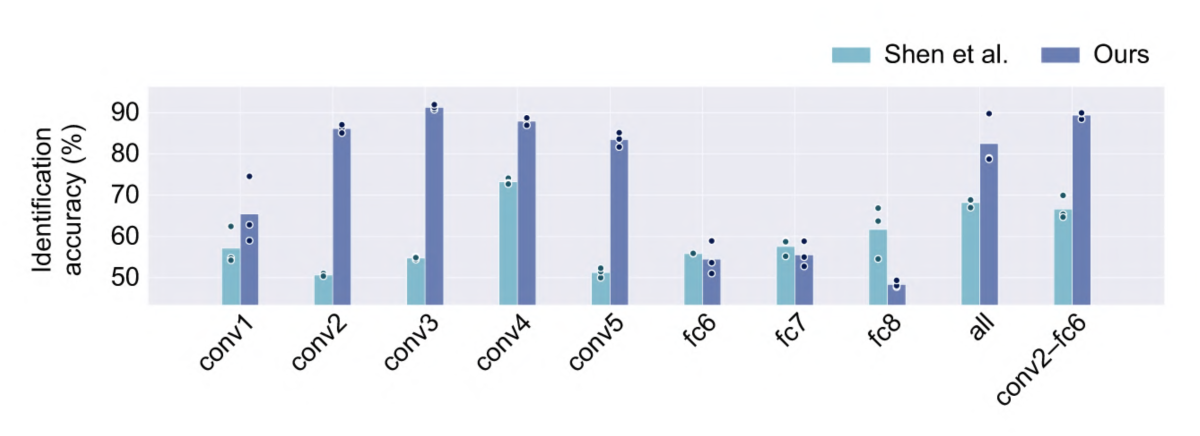
Image identification accuracy for reconstructions of seen artificial shapes. The formats are the same as those in Fig. 2e.

**Supplementary Fig. 8.**
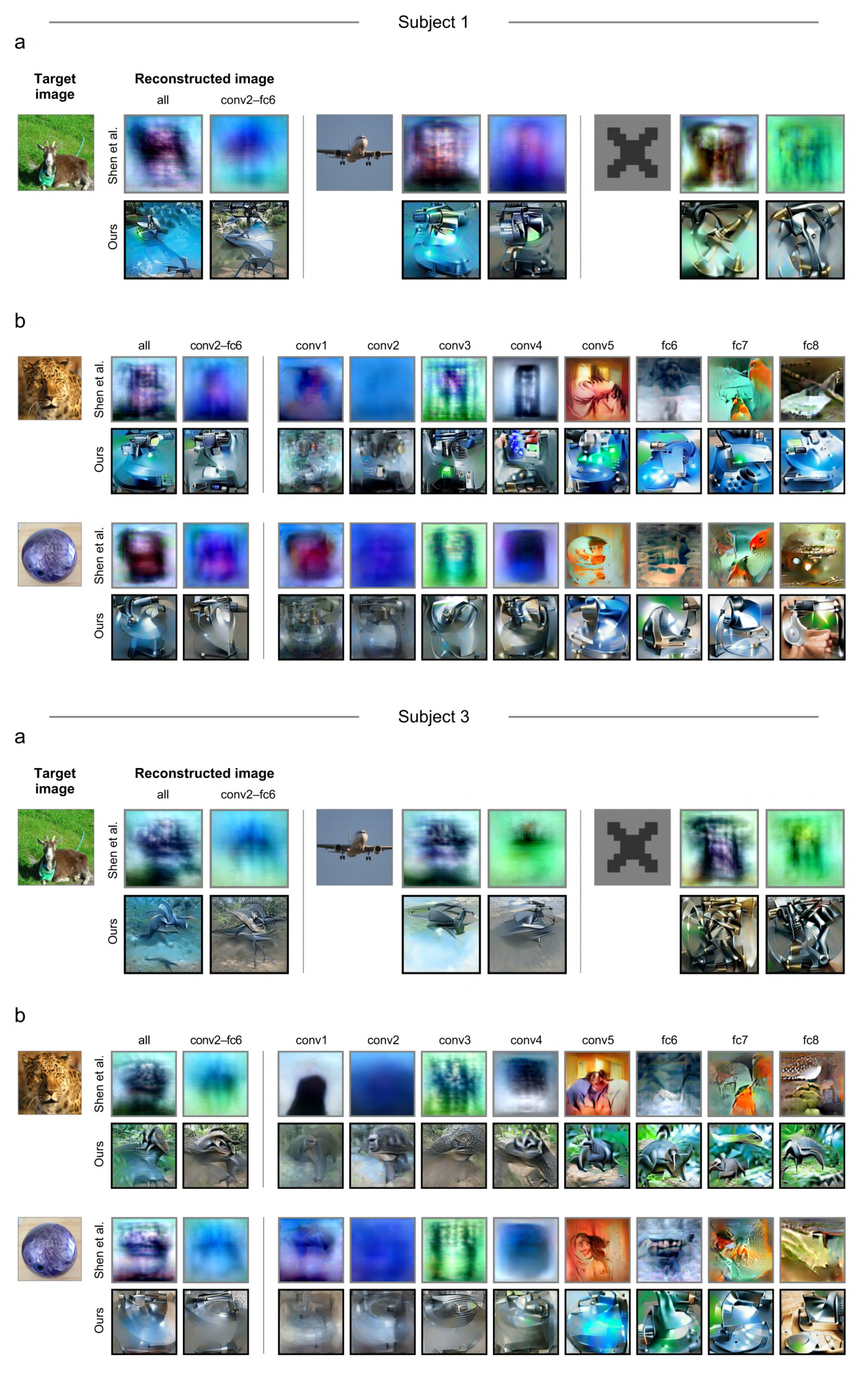
Reconstructions of imagined images from Subjects 1 and 3. The formats and the target images are the same as those in Figs. 3a and 3b.

**Supplementary Fig. 9.**
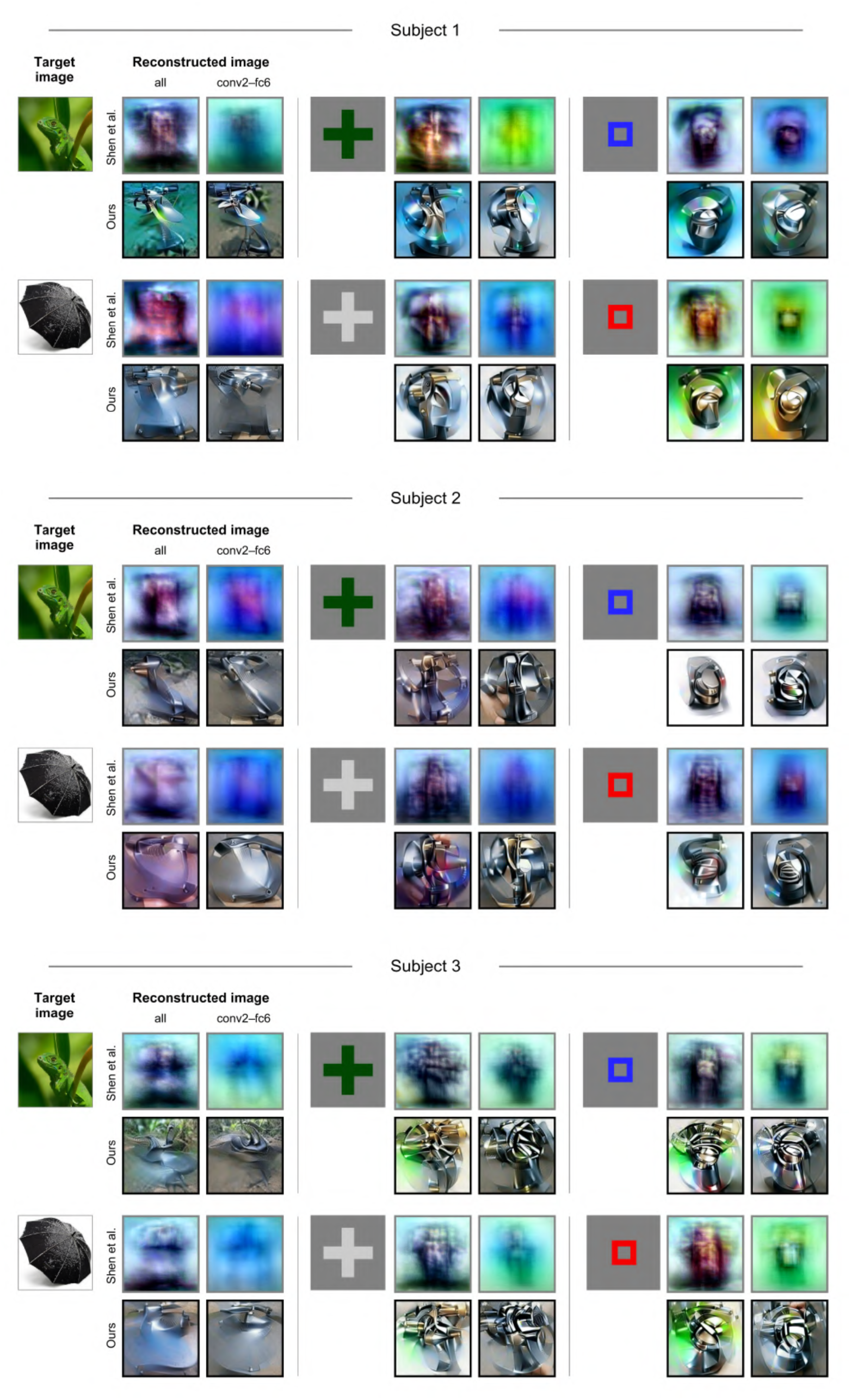
Other examples of imagery reconstructions. The formats are the same as those in Fig. 2a.

**Supplementary Fig. 10.**
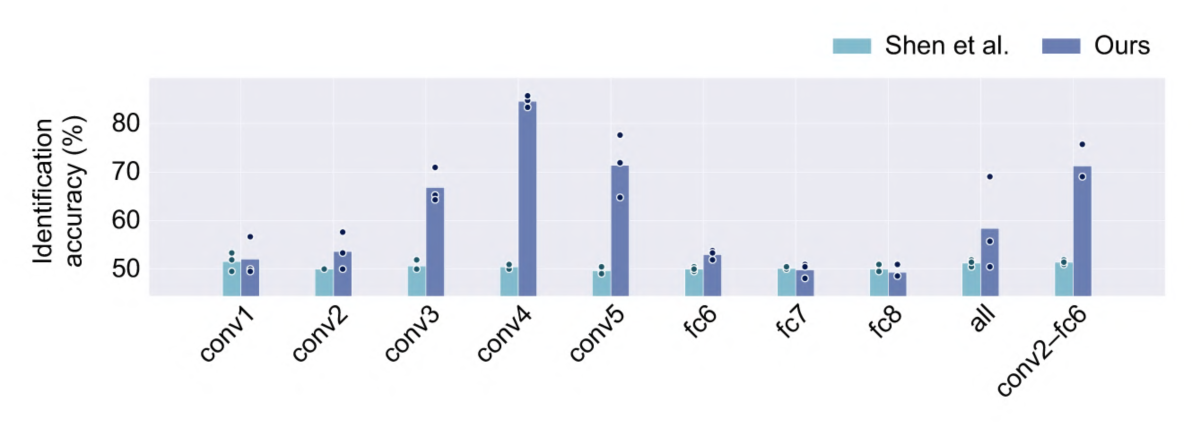
Image identification accuracy for reconstructions of imagined artificial shapes. The formats are the same as those in Fig. 2e.

**Supplementary Fig. 11.**
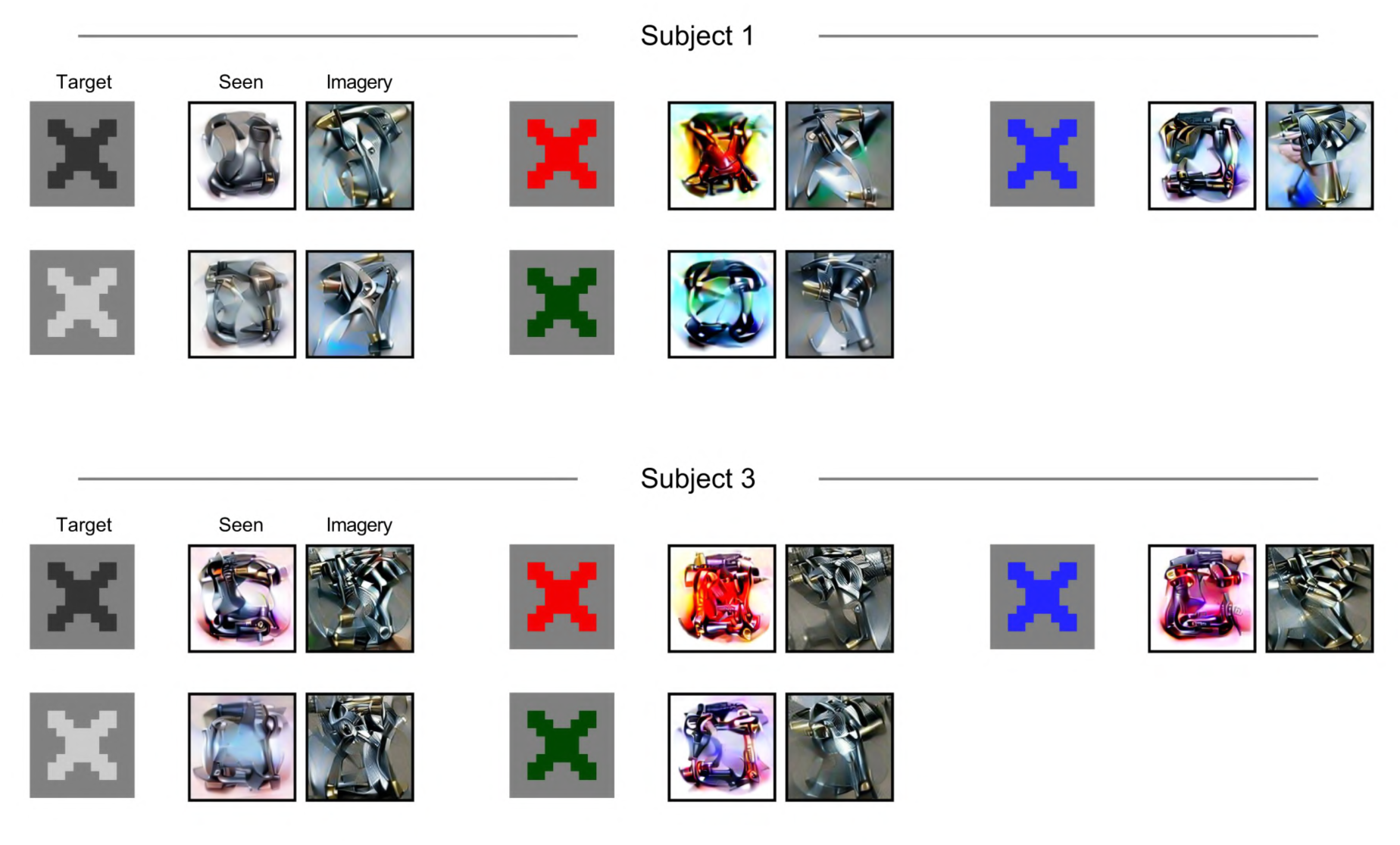
Reconstructions of an X-shaped geometric pattern from Subjects 1 and 3. The formats and target images are the same as those in Fig. 4a.

**Supplementary Fig. 12.**
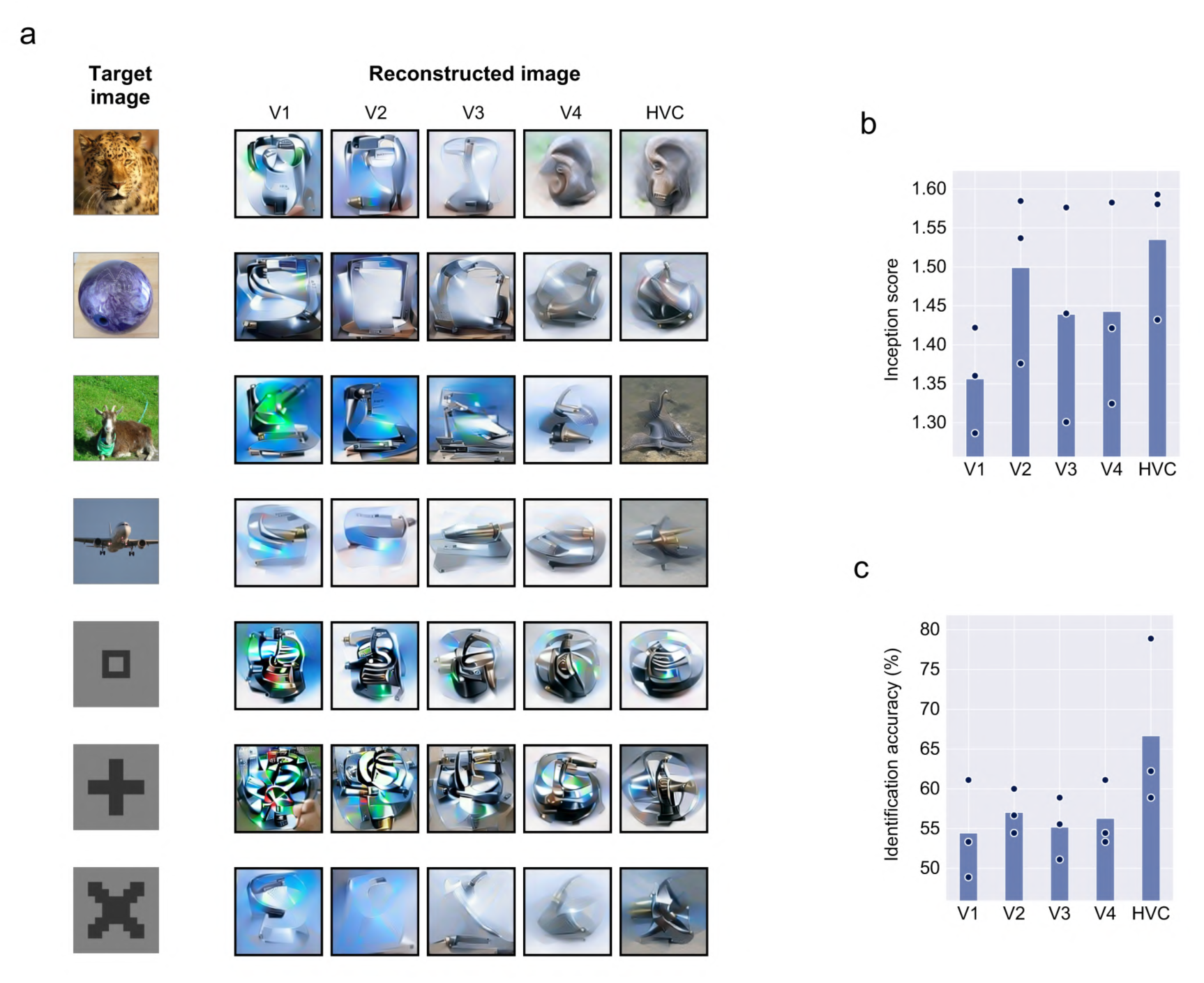
Comparison of imagery reconstructions between input brain areas. (a) Imagery reconstructions from brain areas V1, V2, V3, V4, and the higher visual cortex (HVC). Images reconstructed via conv2–fc6 are shown. **(b)** Inception score. **(c)** Image identification accuracy.

**Supplementary Fig. 13.**
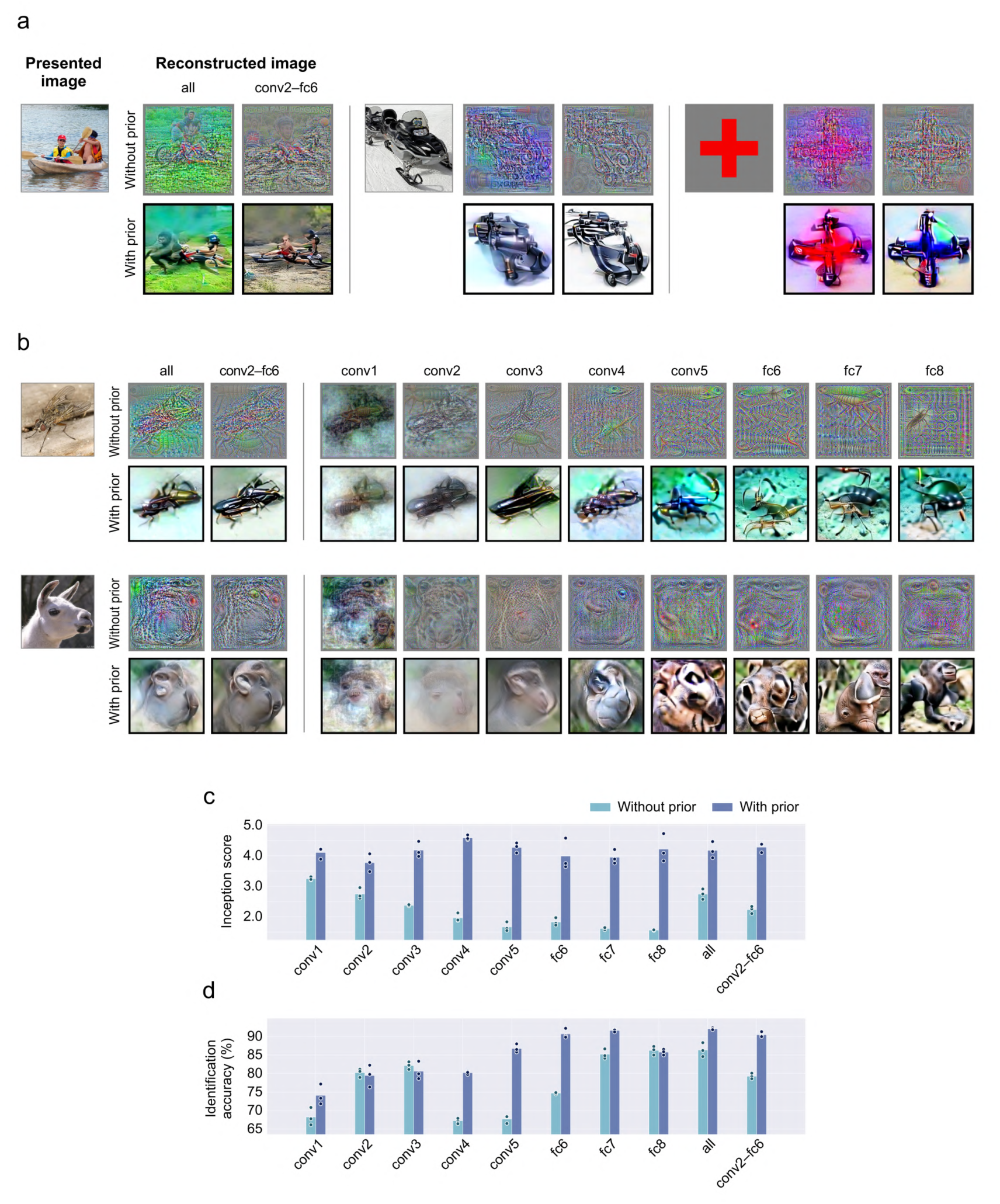
Effect of the image prior (seen image reconstruction). Image reconstruction results are compared between our full method and that without the image prior. The formats and target images are the same as those in Fig. 2.

**Supplementary Fig. 14.**
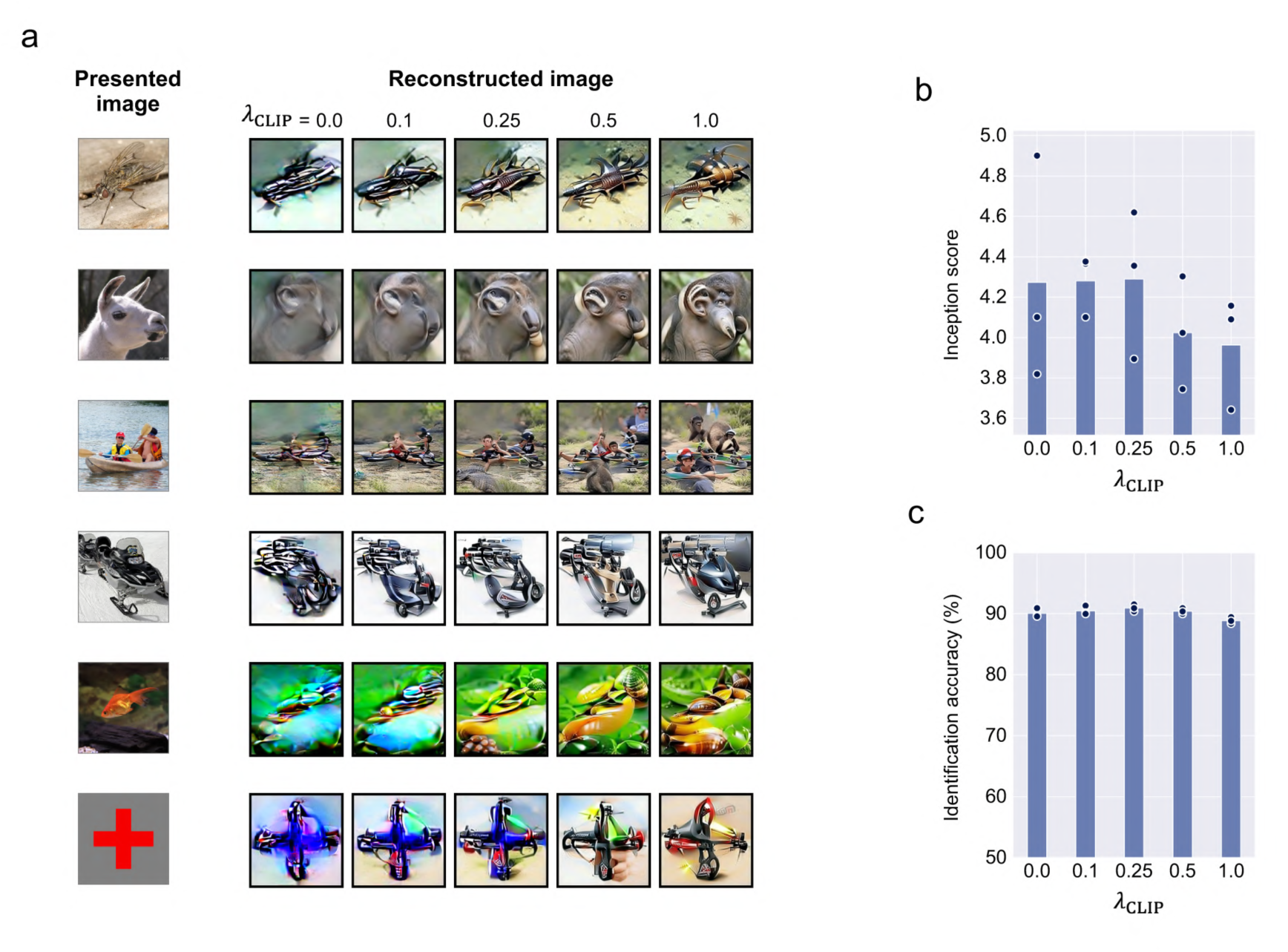
Effect of the assistance of semantic information (seen image reconstruction). Seen image reconstruction results are compared across different strengths of semantic information assistance. The formats are the same as those in Fig. 6.

## References

1. Kay, K. N. & Gallant, J. L. I can see what you see. Nat Neurosci 12, 245 (2009).

2. Rakhimberdina, Z., Jodelet, Q., Liu, X. & Murata, T. Natural Image Reconstruction From fMRI Using Deep Learning: A Survey. Front. Neurosci. 15, 795488 (2021).

3. Cowen, A. S., Chun, M. M. & Kuhl, B. A. Neural portraits of perception: reconstructing face images from evoked brain activity. Neuroimage 94, 12–22 (2014).

4. Lee, H. & Kuhl, B. A. Reconstructing Perceived and Retrieved Faces from Activity Patterns in Lateral Parietal Cortex. J Neurosci 36, 6069–6082 (2016).

5. Schoenmakers, S., Barth, M., Heskes, T. & van Gerven, M. Linear reconstruction of perceived images from human brain activity. Neuroimage 83, 951–961 (2013).

6. Miyawaki, Y. et al. Visual Image Reconstruction from Human Brain Activity using a Combination of Multiscale Local Image Decoders. Neuron 60, 915–929 (2008).

7. Fujiwara, Y., Miyawaki, Y. & Kamitani, Y. Modular Encoding and Decoding Models Derived from Bayesian Canonical Correlation Analysis. Neural Computation 25, 979–1005 (2013).

8. Satake, E., Majima, K., Aoki, S. C. & Kamitani, Y. Sparse Ordinal Logistic Regression and Its Application to Brain Decoding. Front. Neuroinform. 12, 51 (2018).

9. Kay, K. N., Naselaris, T., Prenger, R. J. & Gallant, J. L. Identifying natural images from human brain activity. Nature 452, 352–355 (2008).

10. Naselaris, T., Prenger, R. J., Kay, K. N., Oliver, M. & Gallant, J. L. Bayesian reconstruction of natural images from human brain activity. Neuron 63, 902–915 (2009).

11. Nishimoto, S. et al. Reconstructing visual experiences from brain activity evoked by natural movies. Curr Biol 21, 1641–1646 (2011).

12. Krizhevsky, A., Sutskever, I. & Hinton, G. E. ImageNet Classification with Deep Convolutional Neural Networks. in Advances in Neural Information Processing Systems vol. 25 (Curran Associates, Inc., 2012).

13. Simonyan, K. & Zisserman, A. Very Deep Convolutional Networks for Large-Scale Image Recognition. (2014) doi:10.48550/ARXIV.1409.1556.

14. Radford, A., Metz, L. & Chintala, S. Unsupervised Representation Learning with Deep Convolutional Generative Adversarial Networks. Preprint at http://arxiv.org/abs/1511.06434 (2016).

15. Brock, A., Donahue, J. & Simonyan, K. Large Scale GAN Training for High Fidelity Natural Image Synthesis. Preprint at http://arxiv.org/abs/1809.11096 (2019).

16. Oord, A. van den, Vinyals, O. & Kavukcuoglu, K. Neural Discrete Representation Learning. Preprint at http://arxiv.org/abs/1711.00937 (2018).

17. Razavi, A., Oord, A. van den & Vinyals, O. Generating Diverse High-Fidelity Images with VQ-VAE-2. Preprint at http://arxiv.org/abs/1906.00446 (2019).

18. Esser, P., Rombach, R. & Ommer, B. Taming Transformers for High-Resolution Image Synthesis. in Proceedings of the IEEE/CVF Conference on Computer Vision and Pattern Recognition (CVPR) (2021). doi:10.48550/ARXIV.2012.09841.

19. Song, Y. et al. Score-Based Generative Modeling through Stochastic Differential Equations. Preprint at http://arxiv.org/abs/2011.13456 (2021).

20. Dhariwal, P. & Nichol, A. Diffusion Models Beat GANs on Image Synthesis. Preprint at http://arxiv.org/abs/2105.05233 (2021).

21. Rombach, R., Blattmann, A., Lorenz, D., Esser, P. & Ommer, B. High-Resolution Image Synthesis with Latent Diffusion Models. Preprint at http://arxiv.org/abs/2112.10752 (2022).

22. Seeliger, K., Güçlü, U., Ambrogioni, L., Güçlütürk, Y. & van Gerven, M. A. J. Generative adversarial networks for reconstructing natural images from brain activity. NeuroImage 181, 775–785 (2018).

23. Shen, G., Horikawa, T., Majima, K. & Kamitani, Y. Deep image reconstruction from human brain activity. PLoS Comput Biol 15, e1006633 (2019).

24. Shen, G., Dwivedi, K., Majima, K., Horikawa, T. & Kamitani, Y. End-to-End Deep Image Reconstruction From Human Brain Activity. Front. Comput. Neurosci. 13, 21 (2019).

25. Han, K. et al. Variational autoencoder: An unsupervised model for encoding and decoding fMRI activity in visual cortex. Neuroimage 198, 125–136 (2019).

26. Takagi, Y. & Nishimoto, S. High-resolution image reconstruction with latent diffusion models from human brain activity. http://biorxiv.org/lookup/doi/10.1101/2022.11.18.517004 (2022) doi:10.1101/2022.11.18.517004.

27. Horikawa, T. & Kamitani, Y. Attention modulates neural representation to render reconstructions according to subjective appearance. Commun Biol 5, 34 (2022).

28. Cheng, F., Horikawa, T., Majima, K. & Kamitani, Y. Reconstruction of line illusion from human brain activity. in 2022 Conference on Cognitive Computational Neuroscience (Cognitive Computational Neuroscience, 2022). doi:10.32470/CCN.2022.1149-0.

29. Naselaris, T., Olman, C. A., Stansbury, D. E., Ugurbil, K. & Gallant, J. L. A voxel-wise encoding model for early visual areas decodes mental images of remembered scenes. Neuroimage 105, 215–228 (2015).

30. Harrison, S. A. & Tong, F. Decoding reveals the contents of visual working memory in early visual areas. Nature 458, 632–635 (2009).

31. Albers, A. M., Kok, P., Toni, I., Dijkerman, H. C. & de Lange, F. P. Shared representations for working memory and mental imagery in early visual cortex. Curr Biol 23, 1427–1431 (2013).

32. Xing, Y., Ledgeway, T., McGraw, P. V. & Schluppeck, D. Decoding working memory of stimulus contrast in early visual cortex. J Neurosci 33, 10301–10311 (2013).

33. Horikawa, T. & Kamitani, Y. Generic decoding of seen and imagined objects using hierarchical visual features. Nat Commun 8, 15037 (2017).

34. Reddy, L., Tsuchiya, N. & Serre, T. Reading the mind’s eye: decoding category information during mental imagery. Neuroimage 50, 818–825 (2010).

35. Lee, S.-H., Kravitz, D. J. & Baker, C. I. Disentangling visual imagery and perception of real-world objects. Neuroimage 59, 4064–4073 (2012).

36. Dijkstra, N., Bosch, S. E. & van Gerven, M. A. J. Shared Neural Mechanisms of Visual Perception and Imagery. Trends Cogn Sci 23, 423–434 (2019).

37. Güçlütürk, Y. et al. Reconstructing perceived faces from brain activations with deep adversarial neural decoding. in Advances in Neural Information Processing Systems vol. 30 (Curran Associates, Inc., 2017).

38. Welling, M. & Teh, Y. W. Bayesian Learning via Stochastic Gradient Langevin Dynamics. in Proceedings of the 28th International Conference on International Conference on Machine Learning 8 (2011).

39. Radford, A. et al. Learning Transferable Visual Models From Natural Language Supervision. (2021) doi:10.48550/ARXIV.2103.00020.

40. Deng, J. et al. ImageNet: A large-scale hierarchical image database. in 2009 IEEE Conference on Computer Vision and Pattern Recognition 248–255 (IEEE, 2009). doi:10.1109/CVPR.2009.5206848.

41. Salimans, T. et al. Improved Techniques for Training GANs. Preprint at http://arxiv.org/abs/1606.03498 (2016).

42. Jafari-Khouzani, K. & Soltanian-Zadeh, H. Radon transform orientation estimation for rotation invariant texture analysis. IEEE Trans. Pattern Anal. Mach. Intell. 27, 1004–1008 (2005).

43. Abdelhack, M. & Kamitani, Y. Sharpening of Hierarchical Visual Feature Representations of Blurred Images. eNeuro 5, ENEURO.0443-17.2018 (2018).

44. Nonaka, S., Majima, K., Aoki, S. C. & Kamitani, Y. Brain hierarchy score: Which deep neural networks are hierarchically brain-like? iScience 24, 103013 (2021).

45. Qiao, K. et al. BigGAN-based Bayesian Reconstruction of Natural Images from Human Brain Activity. Neuroscience 444, 92–105 (2020).

46. Crowson, K. et al. VQGAN-CLIP: Open Domain Image Generation and Editing with Natural Language Guidance. in Computer Vision – ECCV 2022 (eds. Avidan, S., Brostow, G., Cissé, M., Farinella, G. M. & Hassner, T.) vol. 13697 88–105 (Springer Nature Switzerland, 2022).

47. Majima, K., Sukhanov, P., Horikawa, T. & Kamitani, Y. Position Information Encoded by Population Activity in Hierarchical Visual Areas. eNeuro 4, ENEURO.0268- 16.2017 (2017).

